# Sequence-specific interactions determine viscoelasticity and aging dynamics of protein condensates

**DOI:** 10.1101/2023.04.06.535902

**Authors:** Ibraheem Alshareedah, Wade M. Borcherds, Samuel R. Cohen, Anurag Singh, Ammon E. Posey, Mina Farag, Anne Bremer, Gregory W. Strout, Dylan T. Tomares, Rohit V. Pappu, Tanja Mittag, Priya R. Banerjee

## Abstract

Biomolecular condensates are viscoelastic materials. Here, we report results from investigations into molecular-scale determinants of sequence-encoded and age-dependent viscoelasticity of condensates formed by prion-like low-complexity domains (PLCDs). The terminally viscous forms of PLCD condensates are Maxwell fluids. Measured viscoelastic moduli of these condensates are reproducible using a Rouse-Zimm model that accounts for the network-like organization engendered by reversible physical crosslinks among PLCDs in the dense phase. Measurements and computations show that the strengths of aromatic inter-sticker interactions determine the sequence-specific amplitudes of elastic and viscous moduli as well as the timescales over which elastic properties dominate. PLCD condensates also undergo physical aging on sequence-specific timescales. This is driven by mutations to spacer residues that weaken the metastability of terminally viscous phases. The aging of PLCD condensates is accompanied by disorder-to-order transitions, leading to the formation of non-fibrillar, beta-sheet-containing, semi-crystalline, terminally elastic, Kelvin-Voigt solids. Our results suggest that sequence grammars, which refer to the identities of stickers versus spacers in PLCDs, have evolved to afford control over the metastabilities of terminally viscous fluid phases of condensates. This selection can, in some cases, render barriers for conversion from metastable fluids to globally stable solids to be insurmountable on functionally relevant timescales.

---

Macromolecular phase transitions give rise to compositionally distinct membraneless bodies in cells that are known as biomolecular condensates ^1, 2, 3^. There is a growing realization that condensates form via a blend of segregative transitions (phase separation) and associative transitions (percolation) ^4^. These processes can give rise to condensates with structural inhomogeneity defined by internal physical crosslinks that impart a network-like organization to condensates ^5, 6, 7, 8, 9^. Structural inhomogeneities and physical crosslinking of molecules give rise to viscoelasticity^10, 11, 12, 13, 14^, whereby network connectivity, combined with the timescales for making and breaking crosslinks and the timescales for molecular transport jointly determine viscoelastic moduli ^15^. The current thinking is that there is an intimate connection between time-dependent viscoelastic moduli of condensates and the functions or pathologies they influence in cells ^16, 17, 18, 19^.

Physical aging of fluid-like condensates is characterized by time-dependent slowdowns of internal dynamics, arrested fusion, and the appearance of irregular morphologies for condensates ^12, 20, 21^. We shall designate the relevant timescale to be *t*_obs_, which is the timescale after condensate preparation when the measurement of interest is performed. For a given value of *t*_obs_, specific measurements are used to probe specific dynamical processes. For a given *t*_obs_, the material properties quantified by measuring dynamical processes can be terminally viscous or elastic depending on the modulus that dominates the long timescale asymptotic behaviors in the measurements. As *t*_obs_ increases, condensates can become dynamically arrested due to high crosslinking density and the mismatch of timescales associated with making and breaking of physical crosslinks versus molecular transport ^7, 22, 23^. This can lead to a decoupling of percolation and phase separation, and a manifestation of percolation (gelation) as arrested phase separation ^7,24, 25, 26, 27^.

Age-dependent dynamical arrest of biomolecular condensates can be described as a glass transition ^12^. In this picture, the terminal material states of condensates at all ages are those of dominantly viscous, albeit dynamically frozen liquids. Glasses are disordered systems and glassy behaviors, which represent a loss of ergodicity ^28^ and mobility of molecules, have been attributed ^12^ to the ruggedness of energy landscapes captured by the random energy model ^29^. The second form of aging is associated with the conversion of disordered fluids to ordered solids ^30, 31, 32^. The morphologies of solid states are typically gleaned from a combination of structural analyses ^20, 26, 33^, mobility measurements, and measurements of increased fluorescence of molecular rotors such as thioflavin T (ThT). The conversion to ordered amyloid fibrils is the current working model for an equilibrium liquid-to-solid description of physical aging.

What are the rules that connect sequence-encoded interactions to viscoelasticities of condensates? Are all physically aged condensates disordered, terminally viscous glasses ^12^, or can aging be a signature of disorder-to-order transitions that generate terminally elastic equilibrium solids? Are the equilibrium solids always amyloid fibers? And what are the sequence-encoded properties that determine the timescales of condensate aging? To answer these questions, we quantified the connections between sequence-encoded interactions and time-dependent material properties of condensates formed *in vitro* by different intrinsically disordered prion-like low complexity domains (PLCDs) ^8, 34, 35^.

The stickers-and-spacers framework ^6^ has been useful for modeling and explaining the origins of sequence-specific driving forces for condensate formation by PLCDs ^8, 34, 35^. Aromatic residues are stickers that form reversible physical crosslinks ^8, 34, 35^. Spacers in PLCDs are primarily Gly, Ser, Asn, and Gln residues. They affect overall solubility while influencing the coupling between phase separation and percolation ^8, 34, 35^. Recent lattice-based coarse-grained simulations reproduced the measured phase behaviors for thirty different variants of A1-LCD ^8^, the PLCD derived from the RNA binding protein hnRNP A1. These computations showed that PLCDs within condensates form physically crosslinked networks with hub-and-spoke-like structures. Multimolecular hubs are defined by the extent of intermolecular physical crosslinking mediated by aromatic stickers, and the spokes are governed by spacer-mediated transport of molecules between hubs, which also requires the breaking of crosslinks ^8, 9^. The interplay between sticker- and spacer-mediated effects leads to spatially inhomogeneous organization of PLCD molecules. Here, our focus is on understanding how network-like structures within PLCD condensates and reconfigurations of these networks determine frequency-dependent viscoelasticity, specifically the storage (elastic) and loss (viscous) moduli ^10, 12, 13, 14, 36, 37^.

## Results

### Measured viscoelastic moduli are governed by network structures of condensates

We employed passive microrheology with optical tweezers (pMOT) ^38^ (**Fig. 1a**) to measure viscoelastic moduli of different PLCD condensates. These experiments yield frequency-dependent storage (*G*′) and loss (*G*″) moduli, which are estimated by analyzing thermal fluctuations of the positions of optically trapped polystyrene particles within condensates (Extended Data Fig. 1a). Passive measurements ensure that the laser power of the trap is kept at a minimal, non-perturbing level ^11^. Frequency-dependent moduli were calculated via Fourier transformation of the position autocorrelation functions of the beads and the optical trap stiffness ^38^ (Extended Data Fig. 1b).

**Fig. 1:**
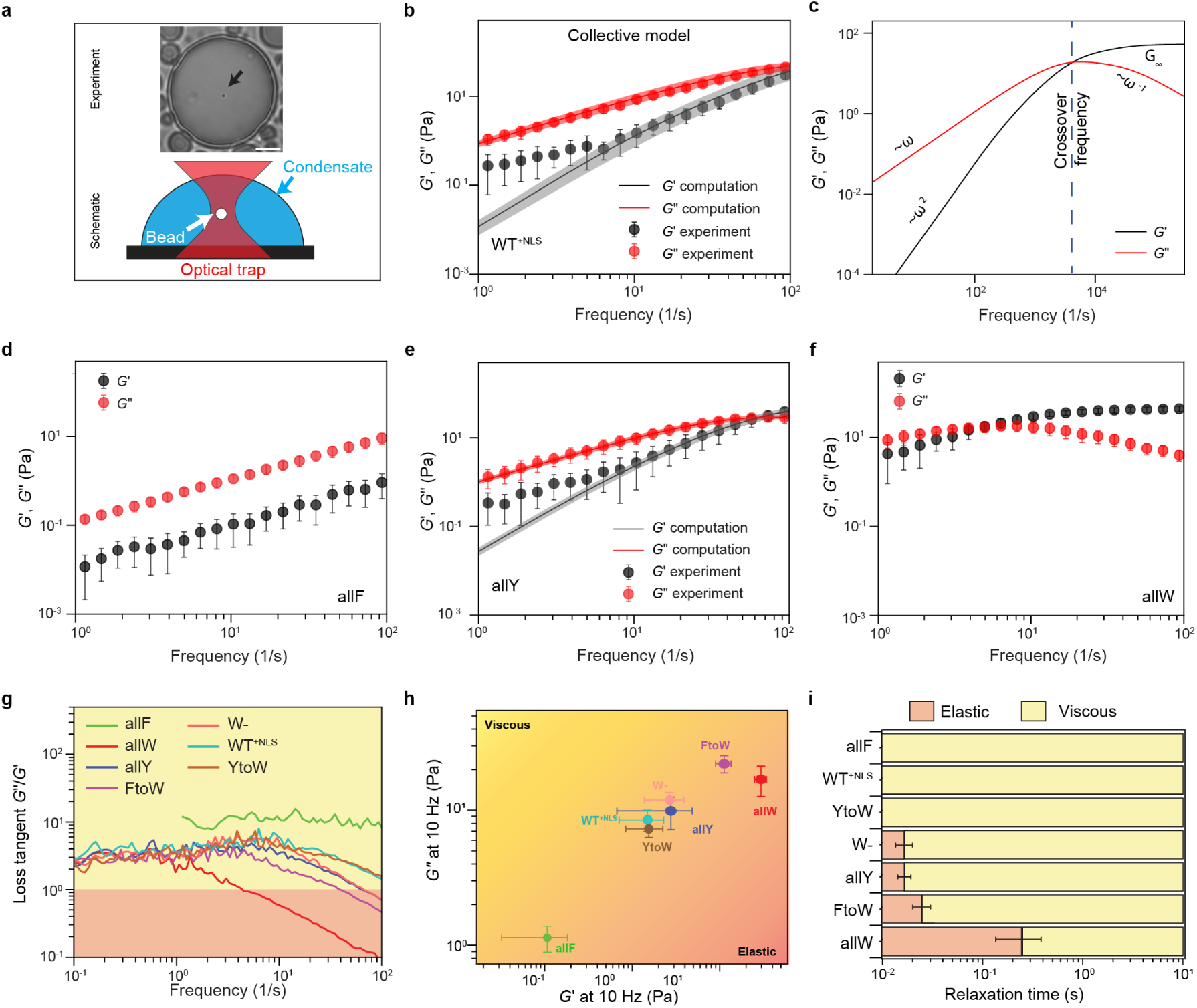
PLCD condensates are viscoelastic Maxwell fluids defined by sequence-specific, frequency-dependent storage (*G*′) and loss (*G*″) moduli. (**a**) Setup for pMOT measurements. The image shows an optically trapped 1 µm polystyrene bead within a condensate formed by WT^+NLS^ A1-LCD. In the schematic, the condensate is shown in blue, the optical trap as a red, hourglass-shaped object, and the bead in white. **(b)** Dynamical moduli for WT^+NLS^ condensates. The figure shows comparisons between measured and computed moduli, where computations use the collective model. **(c)** Dynamical moduli computed from simulations using the collective model for WT^+NLS^ condensates. This highlights the frequency-dependent scaling of storage and loss moduli above and below the crossover frequency, which is annotated by the blue dashed line. *G*′ = *G*″ at the crossover frequency. **(d-f)** Measured dynamical moduli of condensates formed by three sequence variants: **(d)** allF, **(e)** allY, and **(f)** allW. The computed moduli for allF are shown in **Extended Data Fig. 2a**. In panel (e) we see that the collective model provides an accurate accounting of the measured moduli. We lack a coherent LaSSI model for coarse-grained lattice simulations of Trp-containing sequences (see Discussion). For panels (b), and (d)-(f), the error bars for measured data represent standard deviations about the mean that was computed using data for 20 condensates. For panels (b) and (e), the error bands for computed data are standard deviations across 3 replicates. (**g**) Loss tangent plotted against the frequency for condensates formed by seven variants. (**h**) A diagram-of-states based on measured moduli at 10 Hz. (**i**) Delineation of the dominantly viscous versus elastic regimes in terms of relaxation times, computed as the inverse of the crossover frequency, for condensates formed by each of the seven variants.

We designate the wild-type A1-LCD, which also features a nuclear localization signal, as WT^+NLS^. For *t*_obs_ ≈ 5 min, measurements of condensates formed by WT^+NLS^ yield dynamical moduli across a frequency range that spans over two orders of magnitude (**Fig. 1b**). We find that the loss modulus of WT^+NLS^ condensates is an order of magnitude larger than the storage modulus at the lower end of experimental frequencies (0.1-10 Hz), whereas these moduli become equivalent at the crossover timescale of ∼ 10^-2^ seconds. Since the crossover frequency is close to the upper limit of experimentally accessible frequencies in pMOT measurements, we further employed fast video particle tracking (VPT) microrheology to extract mean squared displacements (MSDs) of 200 nm probe particles embedded within WT^+NLS^ condensates. The corresponding MSD curve reveals diffusive motions of the particles at timescales longer than 10 ms (Extended Data Fig. 1c). This is consistent with the pMOT results and suggests that at timescales faster than 10 ms, these condensates are dominantly elastic.

To uncover the sequence-specific connections between condensate structure and measured moduli, we leveraged the results from lattice-based simulations of condensate structures ^8^. Our quantifications of moduli, which are based on simulated structures of condensates, use a graph-theoretic adaptation ^39, 40^ of Rouse-Zimm theory ^41, 42^ for polymer dynamics. The standard implementation of Rouse-Zimm theory is a bead-and-spring model for the dynamics of a single chain ^41, 42^. Key ingredients are: (i) the friction of the medium; (ii) the mobility tensor, including contributions from hydrodynamic interactions; (iii) the Zimm matrix whose elements are determined by the specific interactions of the chain; and (iv) the stochastic contributions due to Brownian motion. Since our focus is on the contributions of equilibrium internal structures of condensates to their viscoelastic moduli, we adopted the simplification of the Rouse-Zimm theory due to Peticolas^43^ and focused exclusively on the eigenvalue spectrum of the Zimm matrix, which is equivalent to a graph Laplacian ^44^. This theory is useful because the solvent is implicit in our simulations, meaning that our Zimm matrices do not account for hydrodynamic interactions.

The spectrum of eigenvalues defines the relaxation times due to interconversions of the network defined by molecules that are physically crosslinked. For a given chain conformation, the diagonal elements of Zimm matrices are the degree, defined as the number of edges incident on a node, where a node is a residue in a chain. The off-diagonal elements specify the connectivity. These are set to –1 if there is an edge between two nodes or 0 if there is no edge. From the published simulations ^8^, we extracted ensembles of condensate-specific graph Laplacians and computed their eigenvalue spectra, which are the relaxation times. The Fourier transform of the superposition of relaxation times shows that the simulated condensates have frequency-dependent storage and loss moduli with a single, system-specific crossover frequency. To put the computed moduli on the same scale as the measurements, we scaled the computed crossover frequency to match experimentally derived values. No other information from the experiment was used in comparing computations to experiments.

Are the dynamics of a single chain in an effective solvent sufficient for explaining measured viscoelastic moduli of PLCD condensates or must one account for the sequence-specific network structures within condensates? We answered this question by comparing measured and computed complex moduli for condensates formed by WT^+NLS^. For the computations, we used two versions of the Zimm matrix namely, the “single-chain model” and the “collective model”. In the “single-chain model”, the nodes represent residues on a single chain within the condensate, and the edges are defined by bonded and non-bonded intra-chain contacts. The Zimm matrix was calculated as *Z = BB^T^* where the unoriented incidence matrix *B* has elements *B_ij_.* For *j* ≠ *i*, the elements *B_ij_* are set to 1 if there is an edge between nodes *i* and *j*; otherwise, *B_ij_* equals zero. For *j=i*, *B_ij_* equals the degree, defined as the number of edges incident on the node *i* ^45^. For the “collective model”, we calculated the Zimm matrices from a graph where the nodes are individual chains, and the edges define any inter-residue contacts between chains. In this model, the collection of molecules in the condensate-spanning network are modeled as a single chain.

The striking observation is that despite not accounting for hydrodynamic effects, the equilibrium ensemble-averaged properties of condensates contain most of the information that is necessary for the faithful reproduction of measured moduli. The only adjustable parameter is the value of the crossover frequency, which we fix using measured values. For condensates formed by WT^+NLS^, the moduli computed using the collective model show very good agreement with the measured moduli (**Fig. 1b**; Extended Data Fig. 1d, 1e). This stands in contrast to the single-chain model (Extended Data Fig. 1f). These results suggest that the moduli, especially the storage modulus, are governed by details of network structures that define how the WT^+NLS^ molecules are organized within the condensate. Our results suggest that viscoelastic moduli are collective properties, defined by the network of intra- and intermolecular interactions. This cannot be captured fully by probing the motions of a single chain ^46^. Instead, the network of molecules becomes an effective single chain in our instantiation of the Rouse-Zimm model that reproduces measured moduli.

### Sticker strengths determine the viscoelastic moduli of PLCD condensates

The measured and computed moduli show a distinct set of features. These are highlighted using the rescaled computed moduli for WT^+NLS^, which enables analysis over a frequency range that spans five orders of magnitude (**Fig. 1c**). There is a single crossover frequency (ω_crossover_) such that *G*′ < *G*″ below ω_crossover_ and *G*′ > *G*″ above ω_crossover_. This points to dominantly viscous behaviors at long timescales whereas elastic behaviors dominate at short timescales. The storage modulus plateaus above ω_crossover_ and varies as ω^2^ below ω_crossover_. The loss modulus varies linearly with ω below ω_crossover_, and as ω^-1^ above ω_crossover_. The presence of a single crossover frequency and the specific functional forms we extract for the frequency dependencies above and below ω_crossover_ imply that WT^+NLS^ condensates probed at *t*_obs_ ≈ 5 min are viscoelastic Maxwell fluids ^12,38^. The apparent deviations from Maxwell-like behaviors at low frequencies (**Fig. 1b**) are likely due to unavoidable noise in the long-time position autocorrelation functions measured using pMOT.

Aromatic stickers follow a hierarchy of strengths whereby Trp is the strongest sticker, Phe is the weakest sticker, and Tyr has strengths that lie between those of Trp and Phe ^8, 10, 22, 35, 47, 48^. Similar distinctions have been observed between purine and pyrimidine moieties in RNA^22, 27^. We designed variants of A1-LCD, where the aromatic stickers were replaced by Phe (allF), Tyr (allY), or Trp (allW) (**Extended Data Fig. 1g**). We also studied variants where only Phe (FtoW), or Tyr (YtoW) residues were replaced by Trp. Finally, in W-, all stickers were Trp residues, and the total number of stickers was reduced by a factor of 0.35 (**Extended Data Fig. 1g**).

Replacing all the aromatic residues in WT^+NLS^ with Phe increases the crossover frequency and lowers the storage and loss moduli (**Fig. 1d**). The increase in ω_crossover_ is substantial, thus preventing its direct measurement using pMOT (**Fig. 1d**). Therefore, we used a numerical method whereby we estimated ω_crossover_ for condensates formed by allF by quantifying it as the value where the curves for *G*′(ω) and *G*″(ω) intersect. Using this method, we estimated ω_crossover_ to be ≈ 4853 Hz (**Extended Data Fig. 2a**). Overall, we observed systematic downshifts of ω_crossover_ as the strengths of stickers increase with the actual values being > 10^2^ Hz for YtoW, 80 Hz for W-, 70 Hz for allY, 60 Hz for FtoW, and 7 Hz for allW (**Fig. 1d-g**, **Extended Data Fig. 2b-2d**). Taken together, we find that ω_crossover_ changes by three orders of magnitude, being the highest for allF and lowest for allW.

The totality of inferences from pMOT measurements for seven variants including WT^+NLS^ can be summarized in three different ways. The loss tangent is a commonly used quantity. It is defined as the ratio of *G*″ to *G*′, which we plot as a function of frequency (**Fig. 1g**). The frequency dependence of this quantity helps identify the timescales for which the condensates are dominantly viscous – loss tangent greater than one – versus timescales for which the condensates are dominantly elastic – loss tangent less than one (**Fig. 1g**). Crossover to predominantly elastic regimes occurs at frequencies that are sequence-specific and encoded by sticker valence and strength.

We can also generate a diagram-of-material-states using variant-specific values of *G*″ and *G*′ at a fixed frequency of 10 Hz (**Fig. 1h**). This analysis shows a one-to-one correspondence between frequency-dependent *G*″ and *G*′ values. The stickers follow a hierarchy whereby Trp is stronger than Tyr, which is stronger than Phe ^8, 10, 22, 35, 47, 48^. As the sticker strengths increase, the values of *G*′ and *G*″ are upshifted, thus showing that the magnitudes of the moduli provide a proxy readout of the strengths of physical crosslinks between stickers. Finally, an alternative depiction of the diagram-of-material-states can be generated to identify the variant-specific relaxation times that correspond to dominantly viscous versus dominantly elastic regimes (**Fig. 1i**). This shows that the timescales corresponding to dominantly elastic behaviors shift toward larger values as sticker strengths increase.

### Bulk viscosities in condensates are inversely correlated with saturation concentrations

Next, we used computations to quantify the temperature dependence of viscosities within PLCD condensates. These viscosities were computed by tracking the timescale associated with the motions of the centers of mass of the molecules within condensates. For all PLCD sequences studied to date, the concentrations of PLCDs in condensates change minimally with temperature except near the critical temperature ^35^. Accordingly, the temperature-dependent saturation concentrations (*c*_sat_) can be used to quantify the free energy change associated with transferring PLCD molecules across the temperature- and variant-specific phase boundary ^35^.

Is there a connection between temperature-dependent intra-condensate viscosities and *c*_sat_ values? We answered this question by quantifying the correlation between computed viscosities and *c*_sat_ values for twenty different PLCD variants across seven different simulation temperatures (**Fig. 2a**). The variants were chosen to ensure that the two-phase regimes span the accessible range of temperatures in the measurements. We observed an inverse correlation between the computed viscosities and *c*_sat_ (**Fig. 2a**). The Pearson *r*-value is negative with a value of –0.996. Strikingly, the computational results for all variants, across all simulation temperatures, collapsed onto a master curve, without requiring any adjustable parameters or fitting. This shows a clear and strong one-to-one correspondence between viscosity and *c*_sat_ and suggests an apparent universality as opposed to system-specificity.

**Fig. 2:**
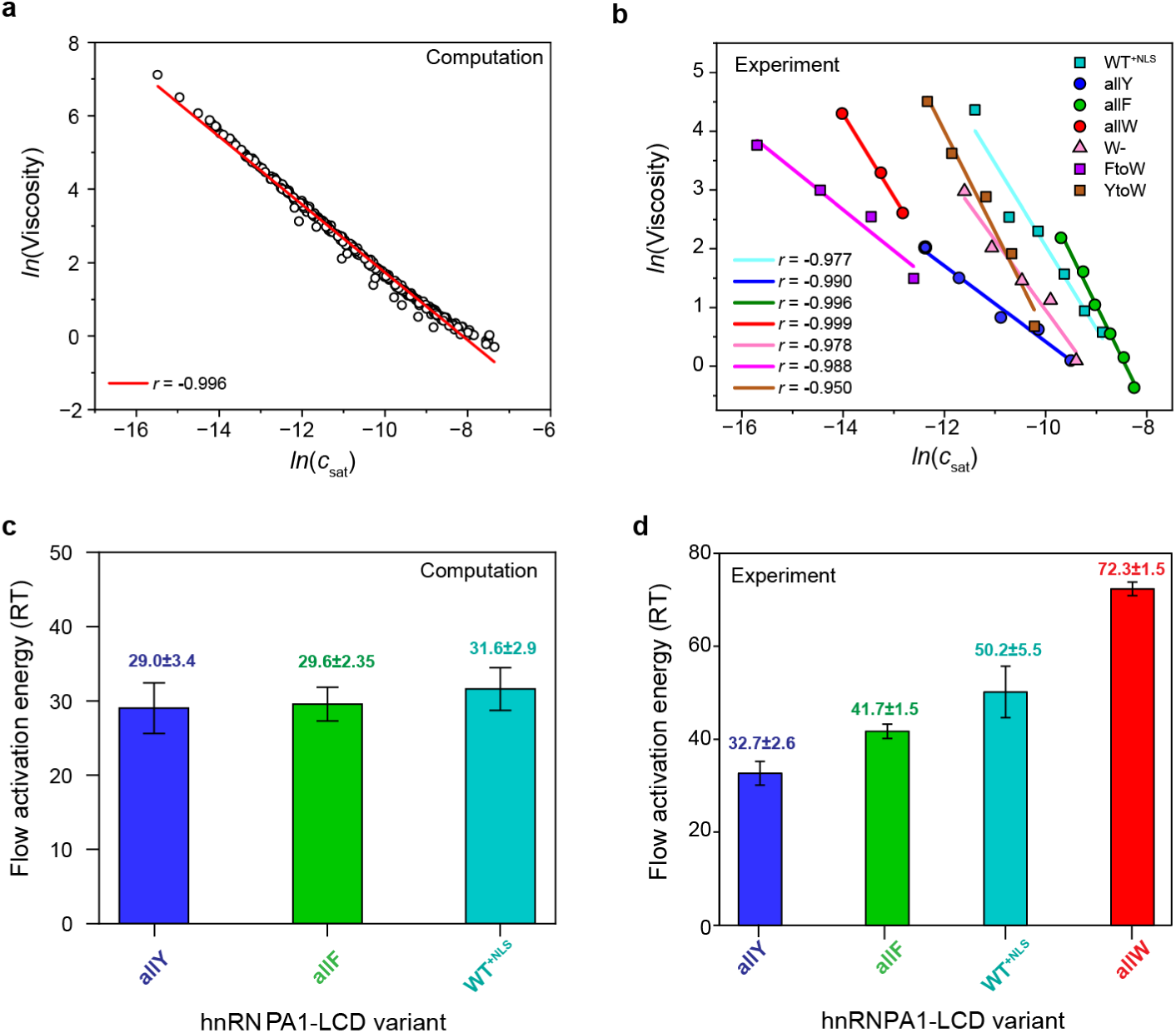
Internal viscosity of condensates and the driving forces for phase separation are inversely correlated. **(a)** Plot of computed, temperature- and variant-dependent viscosity and *c*_sat_ values. The data for different temperatures and variants collapse, without any adjustments, onto a single master curve. The red line is a linear fit to the plot of ln(viscosity) versus ln(*c*_sat_). **(b)** Plots of the measured temperature-dependent viscosities versus *c*_sat_ for seven different systems (see legend). However, while there is a strong negative correlation between ln(viscosity) versus ln(*c*_sat_), the slopes are variant-specific. (**c**) Computed flow activation energies for allY, allF and WT^+NLS^. (**d**) Measured flow activation energies for allY, allF, WT^+NLS^, and allW. Error bars in (**c**) and (**d**) represent the error obtained from the linear fit.

We tested the robustness of the computational results by quantifying the strength of the correlation between dynamical bulk viscosity and *c*_sat_. We measured mean squared displacements (MSDs) of probe particles within PLCD condensates using video particle tracking (VPT) as a function of temperature (**Extended Data Fig. 3a**). The VPT measurements allow us to monitor the motions of 200 nm probe particles in condensates for longer timescales (hundreds of seconds) and over longer distances that are not constrained by the optical trap. This enabled us to measure viscosities at longer timescales, with improved statistics afforded by being able to track tens to hundreds of independent probe particles (**Extended Data Fig. 3b**). For each of the seven variants, the measurements showed clear negative correlations between viscosity and *c*_sat_ (**Fig. 2b**). The Pearson *r*-values are akin to those we estimate from computations. However, unlike the computations, we now observe that plots of measured viscosity versus *c*_sat_ do not collapse onto a single master curve. This cannot be due to discrepancies between computed and measured *c*_sat_ values, because these are essentially identical to one another ^8^. Instead, the variant-specific slopes and intercepts in the measurements point to intra-condensate hydrodynamic effects, which are missing in the simulations, and are likely the result of correlations between solvent-mediated, system-specific interactions in condensates.

The differences between computations and measurements allow us to quantify the magnitudes of proposed hydrodynamic contributions using a method of comparative slopes. Here, we denote *m*_0_ to be the slope of the master curve derived from the plot of ln(viscosity) versus ln(*c*_sat_) using computations. Next, we denote *m_i_* as the slope for variant *i* derived using system-specific measurements. The ratio *s_i_* = (*m_i_* / *m*_0_) quantifies the contribution from hydrodynamics for variant *i*. A value of *s_i_* that is less than unity implies that hydrodynamic interactions weaken the correspondence between dynamical properties of condensates and the free energy of transfer between the dilute and dense phases. A value of *s_i_* that is greater than unity implies that a favorable transfer free energy from dilute to dense phases (lower *c*_sat_ values) is amplified by intra-condensate hydrodynamic interactions. From comparisons of measurements and computations, we find that the *s_i_* values are 0.7, 1.5, and 1.9, for allY, allF, and WT^+NLS^, respectively. The implication is that hydrodynamic and thermodynamic effects show negative cooperativity for allY and positive cooperativity for allF and WT^+NLS^. These effects do not follow the hierarchy of sticker-sticker interactions. Instead, they appear to be governed by a system-specific interplay of solvent-chain, and chain-chain interactions within condensates whereby variants with a higher relative Phe or Trp contents show steeper slopes, while variants with higher Tyr contents exhibit shallower slopes.

For a given temperature, viscosities quantify the frictional effects that arise due to the molecular and mesoscale interactions among molecules that are solvent-mediated. The temperature dependence of viscosity quantifies the energy barrier to reconfiguration of the network of crosslinked molecules. This can be extracted using thermorheology ^49, 50^. These barriers are referred to as flow activation energies because they represent the thermal energy required to overcome the barrier to enable the flow of molecules. To quantify flow activation energies, we measured MSDs of probe particles within PLCD condensates using VPT as a function of temperature. This allowed us to derive temperature dependencies of viscosities of different condensates (**Extended Data Figs. 3c, 3d**). The measured viscosities show an exponential dependence on temperature that is consistent with an Arrhenius relationship (**Extended Data Fig. 3e**) ^50, 51^. Similar results were obtained using computations (**Extended Data Fig. 3f**). The Arrhenius relationship highlights the presence of a single, system-specific barrier to network reconfiguration.

From fits to the Arrhenius relationships, we extracted flow activation energies using data from measurements and computations in units of *RT*, where *T* = 25°C and *R* is the ideal gas constant. We compared the magnitudes of the flow activation energies derived from computations (**Fig. 2c** and **Extended Data Fig. 3g**) to those derived from VPT measurements (**Fig. 2d**). To enable the comparisons, we denote the computed barrier for condensates formed by sequence *i* as *E_F_*_,0*i*_. The computed values do not account for hydrodynamic effects, and hence they can only quantify the contributions of network structures that form within condensates. The values we obtain are sequence-specific, although they are narrowly distributed between 26 *RT* and 32 *RT* (**Fig. 2c**, **Extended Data Fig. 3g**). The values we obtain for *E_F_*_,0*i*_ reflect the similar small-world network topologies and the sequence-specificity of extent of crosslinking within simulated condensates ^8^. As shown previously, the extent of crosslinking does not correlate strongly with *c*_sat_. This is true of the computed values of *E_F_*_,0*i*_ as well.

Next, denoting *E_F_*_,*i*_ (**Fig. 2d**) as the measured flow activation energy for system *i*, we computed the ratio *E_F_*′ = (*E_F_*_,*i*_ / *E_F_*_,0*i*_). This quantifies system-specific hydrodynamic contributions to the flow activation energy that are not present in the lattice simulations. If *E_F_*′ equals unity, there are negligible contributions from hydrodynamic effects to the energy barrier for network reconfiguration. Conversely, if *E_F_*′ is less than unity, then hydrodynamic effects lower the barrier, and the opposite is true if *E_F_*′ is greater than unity. From comparisons of measurements and computations, we find that the *E*_F_′ values are ≈ 1.1, 1.4, and 1.6 for allY, allF, and WT^+NLS^, respectively. The implication is that hydrodynamic effects play a minor role in determining the flow activation barrier for allY, and they increase the barriers for allF and WT^+NLS^. Taken together with the comparative slope analysis, we find that there is negative cooperativity between hydrodynamic effects and the strengths of intermolecular interactions in determining the viscosity of allY condensates. However, the barrier to flow activation is negligibly affected by hydrodynamic effects. For the allF and WT^+NLS^ condensates, there is positive cooperativity between hydrodynamic effects and the strengths of intermolecular interactions as determinants of bulk condensate viscosity, and an enhancement of the barrier to network reconfiguration by hydrodynamic effects. Taken together, our results highlight an intimate, and hitherto unappreciated interplay between molecular grammars and hydrodynamic effects that break apparently universal relationships, doing so in system-specific manners.

### Mutations to spacers that weaken the drive for condensation can drive dynamical arrest

Previous studies showed that the replacement of Gly residues with Ser weakens the driving forces for condensation of PLCDs ^35^. This effect could be explained by the higher effective solvation volume of Ser over Gly ^52^. However, even though substitutions of Gly to Ser weakened the driving forces for condensation, simulations showed that the extent of crosslinking within condensates remained on a par with or was even higher than what was observed for condensates formed by WT and allY ^8^. These observations point to a separation of functions between networking and density transitions that can be unmasked via mutations to spacers ^8^. The extent of internal crosslinking within condensates, which is governed by the associativity of multivalent macromolecules ^4^, does not have to follow the driving forces for phase separation, which are segregative transitions driven by solubility considerations. To test the proposal of separation of functions, we measured the dynamical properties of condensates formed by variants of A1-LCD where different numbers of Gly residues were replaced with Ser **(Extended Data Fig. 4a).**

VPT measurements and computations showed that in allY, allY^20GtoS^, and allY^30GtoS^ condensates studied at *t*_obs_ ≈ 5 min, the bulk viscosities of condensates decrease with weakened driving forces for phase separation (**Fig. 3a**). Condensates formed by the variant WT^30GtoS^ are highly dynamic at early time points, as seen by the motion of embedded 200 nm polystyrene particles. However, they showed evidence of dynamical arrest within *t*_obs_ < 10 min (Movie S1, Extended Data Fig. 4b). We quantified the average diffusion coefficients of multiple probe particles within WT^30GtoS^ condensates as a function of *t*_obs_. The average diffusion coefficient decays to zero rapidly with increasing *t*_obs_ (**Fig. 3b**). This indicates that Gly-to-Ser mutations accelerate the aging behavior compared to WT^+NLS^ condensates (Extended Data Fig. 4b). This accelerated aging prevented quantitative measurements of viscosity and viscoelastic properties of WT^30GtoS^ condensates. However, introducing Gly-to-Ser substitutions into the allY variant (allY^30GtoS^) leads to a decay of the average diffusion coefficient of the probe particles to zero within *t*_obs_ ∼1.5 hrs (**Fig. 3c**). Importantly, we were able to tune the timescales for the onset of dynamical arrest in the allY system by titrating the Gly-to-Ser substitutions. This is shown in root mean squared displacement (RMSD) histograms collected from particle tracking measurements of more than 10^3^ probe particles within different condensates for *t*_obs_ < 1 hrs and at *t*_obs_ ≈ 3 hrs or 24 hrs (**Fig. 3d**, top, middle, and bottom).

**Fig. 3:**
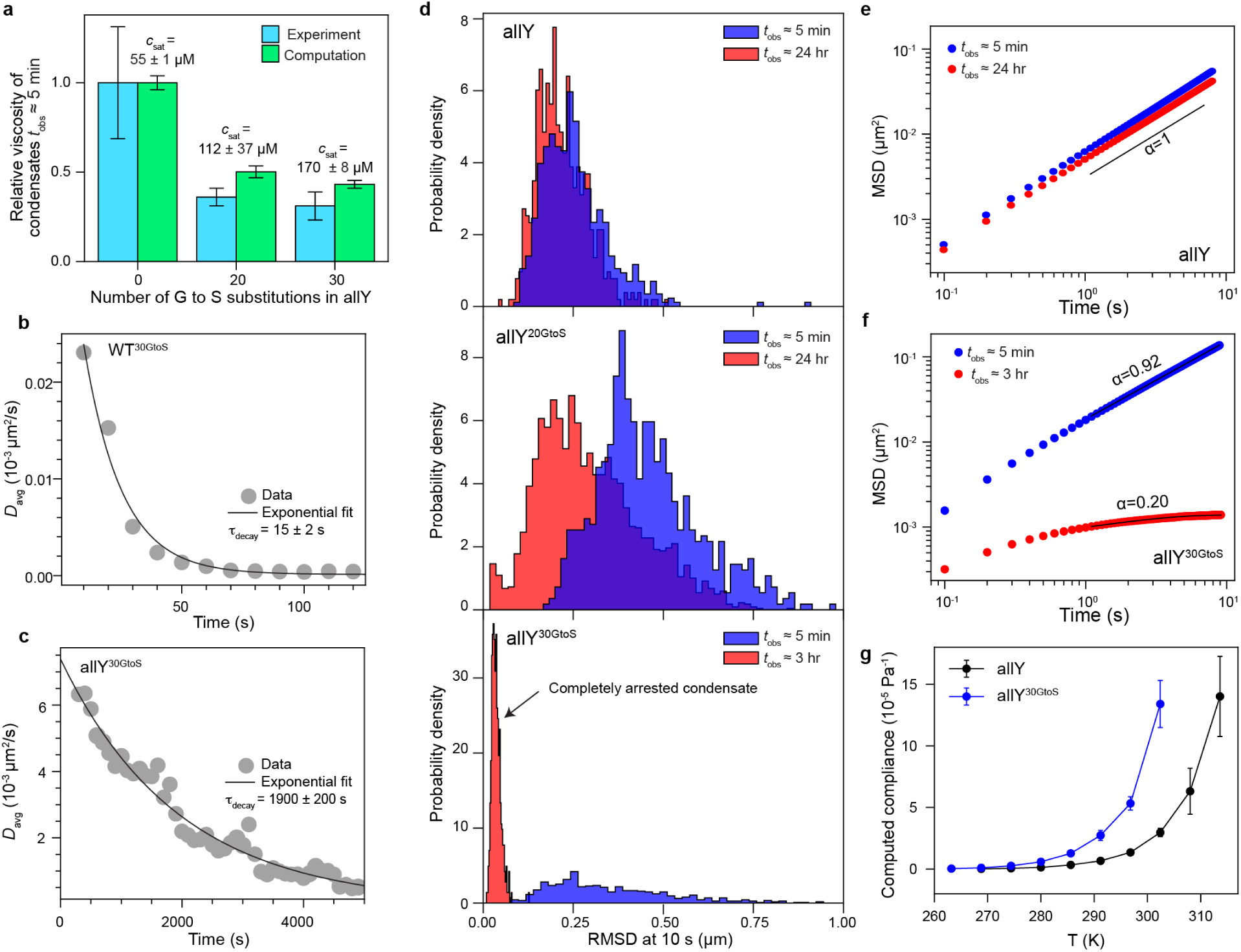
Mutations that alter spacers of A1-LCD accelerate the dynamical arrest of condensates. **(a)** Relative changes in viscosities within condensates at *t*_obs_ ≈ 5 min, calibrated against the allY system, and quantified as a function of different numbers of Gly to Ser substitutions. Data are shown for relative viscosities from measurements and computations, where the latter uses the collective model for the Zimm matrix. Error bars represent the standard deviation in the estimate of the mean relative viscosity. Experimental viscosities are obtained from at least 5 experiments featuring at least 50 tracked particles in each measurement. *c*_sat_ values shown for ≈ 20 °C. **(b)** Average diffusion coefficient of 200 nm polystyrene beads within WT^30GtoS^ condensates measured as a function of time following the start of sample imaging. The black line shows an exponential decay fit with the characteristic decay times τ_decay_ as indicated. **(c)** Same as **(b)** but for allY^30GtoS^ condensates. **(d)** Histograms for the Root Mean Squared Displacement (RMSD) of 200 nm polystyrene beads measured at 10 s lag times within condensates at *t*_obs_ ≈ 5 min and *t*_obs_ ≈ 24 hrs. The top, middle, and bottom panels show histograms for condensates at different times of *t*_obs_ (ages of condensates are in the legends) formed by allY, allY^20GtoS^, allY^30GtoS^, respectively. **(e)** Ensemble-averaged MSD of 200 nm beads in allY condensates at *t*_obs_ ≈ 5 min (blue) and *t*_obs_ ≈ 24 hrs (red). The solid black line is the fit to the equation *MSD*(*τ*) = 4*Dτ^α^* + *N*. The value of the exponent obtained from the fit shows diffusive motion. **(f)** Same as **(e)** but for allY^30GtoS^ condensates. The values of the exponents show the onset of sub-diffusive motion **(g)** Temperature-dependent compliance computed from the simulations for condensates formed by allY and allY^30GtoS^. We use the collective model for computing the Zimm matrix. Error bars are standard deviations of the mean compliance values computed using 3 replicates.

In the absence of spacer mutations, the ensemble-averaged MSDs of probe particles within allY condensates did not show any significant alterations even after 24 hrs (**Fig. 3e**). In contrast, the MSD measurements showed age-dependent dynamics for condensates formed by the allY^20GtoS^ and allY^30GtoS^ variants (Extended Data Fig. 4c, d, and **Fig. 3f**). For *t*_obs_ ≈ 5 min, the particles within allY^30GtoS^ condensates showed diffusive behavior as expected for a material that behaves as a viscous fluid at long timescales. However, after *t*_obs_ ≈ 3 hr, the observed MSD curves were sublinear (**Fig. 3f**), indicating sub-diffusive motion of the probe particles. The flatness of the MSD curves at longer lag-times indicates that the translational diffusion of probe particles becomes caged due to the onset of stable elastic networks ^20, 53, 54, 55^ in condensates formed by allY^30GtoS^. Indeed, the histogram of RMSDs, which initially showed a peak around an RMSD of ∼0.25 µm, showed a sharp peak near zero within 3 hours, indicating complete dynamical arrest of the condensate (**Fig. 3d**, bottom).

Taken together, the picture that emerges is as follows: Gly-to-Ser substitutions on a common allY background of stickers weaken the driving forces for condensate formation. Viscosities of condensates observed at *t*_obs_ ≈ 5 min are lowered with increased number of Gly-to-Ser substitutions. However, because the sticker valence remains the same as in the allY system, the extent of crosslinking within condensates remains high. Therefore, the Gly-to-Ser substitutions show a combination of high early-time mobility and high degrees of crosslinking. This combination leads to unexpected behaviors such as the onset of dynamical arrest and accelerated physical aging of the dense phase. Notably, the timescales for complete dynamical arrest decrease with increasing numbers of Gly to Ser substitutions, highlighting the programmability of dynamical arrest through mutations to spacers (**Fig. 3d**).

What may be the physical origin of accelerated dynamical arrest caused by Gly-to-Ser substitutions? Increasing the number of Gly-to-Ser substitutions increases the compliance of networks formed within condensates that are dominantly viscous (**Fig. 3g**). This was inferred from simulation results. The zero-shear compliance quantifies the maximal network reconfiguration achievable at a given stress. Across all temperatures, the compliance was found to be systematically higher for variants with increased numbers of Gly-to-Ser substitutions (**Fig. 3g**). Increased compliance should enable facile network rearrangements. However, these facile network rearrangements, which happen in a material with a high degree of internal crosslinking, appear to engender a mismatch in timescales whereby rapid mobilities are hindered by the high density of crosslinks, thus leading to dynamical arrest ^7^ and physical aging. If this prediction is valid, then the allY^30GtoS^ condensates should become dominantly elastic as *t*_obs_ increases. We tested this prediction by calculating viscoelastic properties as a function of *t*_obs_ via analysis of MSDs of the probe particles (see Supplementary Material) ^56, 57^. Upon converting the measured values of MSDs to complex moduli, we find that allY^30GtoS^ condensates are primarily viscous at *t*_obs_ ≈ 5 min (Extended Data Fig. 5a-5c). Specifically, the loss modulus is an order of magnitude higher than the storage modulus within the experimental frequency range (∼0.1 to 10 Hz). The elastic modulus is on the order of ∼0.01 Pa, and its rapid fluctuation reflects the noise of the MSD measurement at long lag times where the time-averaged statistics are much lower than for short lag times. Importantly, we find that, upon physical aging, the elastic modulus increases dramatically for condensates of allY^30GtoS^, which are dynamically arrested at *t*_obs_ ≈ 3 hrs. The overall shapes of the frequency-dependent viscoelastic moduli resemble that of Kelvin-Voigt solids (Extended Data Fig. 5d-f) and confirm that allY^30GtoS^ condensates are dominantly elastic at long observation times (*t*_obs_ ≥ 3 hrs).

### Active microrheology affirms the terminally elastic nature of aging condensates

To further investigate the nature of the dynamical transitions that accompany condensate aging upon mutations of spacer residues, we employed active microrheology using an optical tweezer-based creep test ^58^ (**Fig. 4a** and **Extended Data Fig. 6a**). In the creep tests, a polystyrene particle is trapped within the condensate and dragged through the condensate at a constant velocity (**Fig. 4a**). This ensures a constant strain rate within the condensate. We recorded how the particle follows the traveling optical trap, and simultaneously measured the forces exerted on the particle by the internal networks of condensates. The expected response of a material with terminally viscous behavior is of a particle that follows the trap until the optical trap is stopped, and remains at the new position once the trap is stopped ^59^ (**Fig. 4b, top** and **Extended Data Fig. 6a, bottom**). For a terminally viscous Maxwell fluid, the bead will also follow the trap but with a lag time that is governed by the terminal relaxation time of the fluid ^59^ (**Fig. 4b, middle** and **Extended Data Fig. 6a, middle**). Finally, for a terminally elastic material such as a Kelvin-Voigt viscoelastic solid, the particle will follow the trap to a certain extent before recoiling to its initial position ^59^ (**Fig. 4b, bottom** and **Extended Data Fig. 6a, top**). Local motions of the particle at short timescales are allowed whereas, at longer timescales, the particle is constrained within physical cages that are set up by elastic networks that form on system-specific length scales ^60^.

**Fig. 4:**
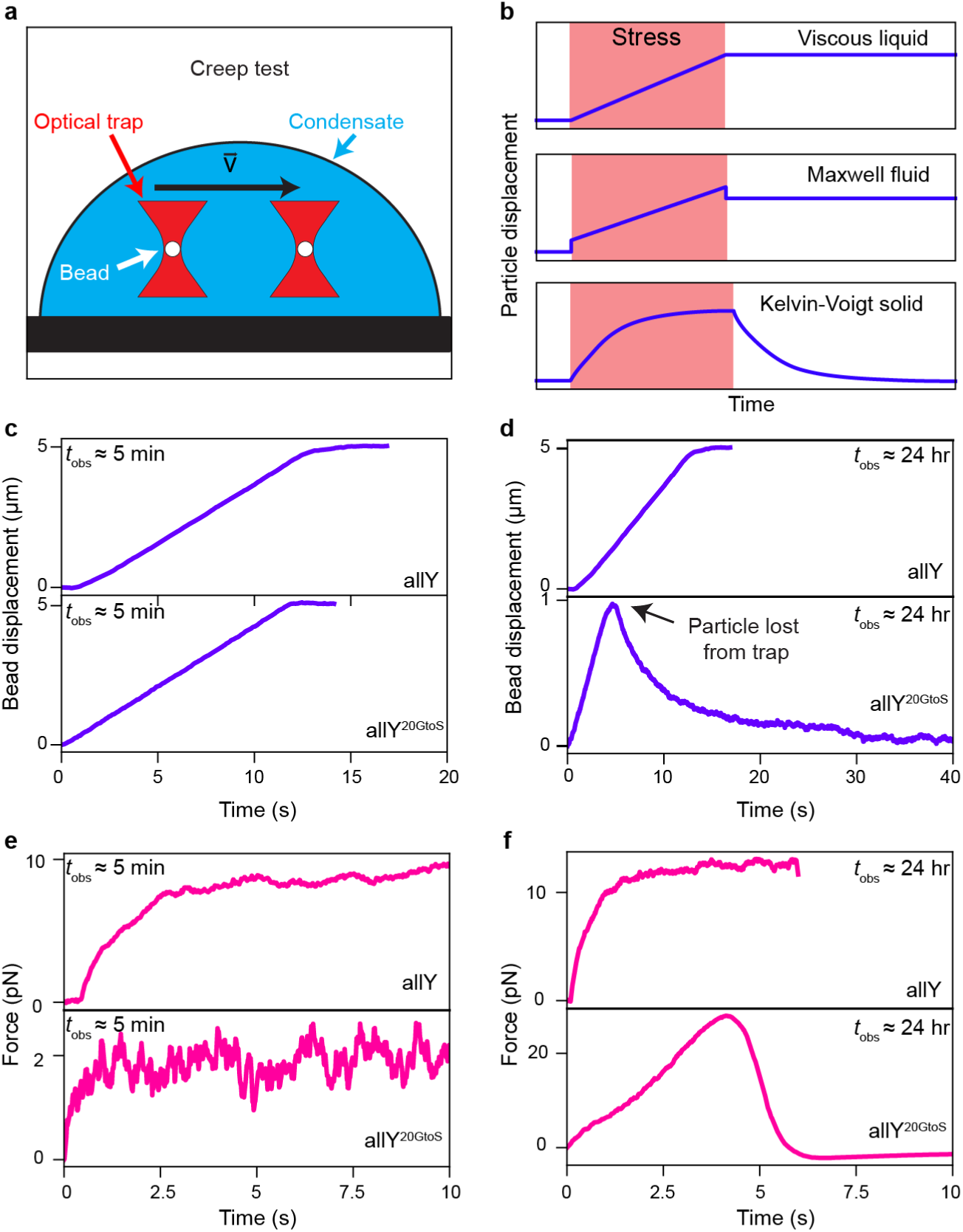
Dynamically arrested PLCD condensates resemble Kelvin-Voigt solids with terminally elastic behavior. **(a)** Schematic of the optical tweezer-based creep test microrheology experiment. A bead is trapped within a condensate using an optical trap. The trap is programmed to move a set distance (here 5 µm) at a constant velocity (0.5 µm/s). The bead position and the force exerted by the condensate on the bead are monitored via a camera and a quadrant photodetector, respectively. **(b)** Expected bead position in response to the trap movement during the creep test for a viscous liquid, a viscoelastic Maxwell fluid, and a Kelvin-Voigt solid. **(c)** Bead position as a function of time in response to the programmed trap movement in allY and allY^20GtoS^ condensates at *t*_obs_ ≈ 5 min. **(d)** Same as (**c**) but for aged allY and aged allY^20GtoS^ condensates (*t*_obs_ ≈ 24 hrs). Note that the bead in allY^20GtoS^ condensate recoils back to its initial position after ∼1 µm travel, indicating that the condensate is exerting an elastic force that overpowers the force exerted by the optical trap. As shown in panel **(b**), this response is characteristic of a viscoelastic solid. **(e)** Force exerted on the bead by the condensate network during the creep test experiment. Data are shown for allY and allY^20GtoS^ condensates at *t*_obs_ ≈ 5 min. The saturation of the force is a characteristic of the viscous drag force, which depends on the velocity of the trap, the viscosity of the condensate, and the size of the bead. **(f)** Same as **(e)** but for condensates at *t*_obs_ ≈ 24 hr. The data in the bottom of panel **(f)** indicate that aged condensates formed by allY^20GtoS^ behave like Kelvin-Voigt solids. Comparison with the top panel highlights the fact that these condensates transition from being Maxwell fluids at early time points to elastic, Kelvin-Voigt solids as they age.

We performed optical tweezer-based creep tests on allY and allY^20GtoS^ condensates. At *t*_obs_ ≈ 5 min, both condensates showed terminally viscous fluid behavior where the bead traveled along with the optical trap for the entire traveling distance of 5 µm (**Fig. 4c**). At *t*_obs_ ≈ 24 hrs, the allY condensates still displayed terminally viscous behaviors (**Movie S2,** **Fig. 4d, top**). However, the allY^20GtoS^ condensates exhibited behaviors consistent with viscoelastic solids (**Movie S3,** **Fig. 4d, bottom**). Specifically, the probe particle was lost from the optical trap at ∼1 μm. Upon being lost from the trap, the probe particle recoiled back to its original position as expected for a Kelvin-Voigt solid.

We also measured the forces exerted on the probe particle by condensate networks during creep test experiments. In nascent allY and allY^20GtoS^ condensates, the saturation of the force is a characteristic of the viscous drag force ^61^, which depends on the velocity of the trap, and the viscosity of the condensate, but is independent of the distance (**Fig. 4e, top**). While this behavior is preserved for the allY condensates investigated at *t*_obs_ ≈ 24 hrs (**Fig. 4e, bottom**), the profile is very different for the aged condensates formed by allY^20GtoS^ (**Fig. 4f, top** versus **bottom**). We systematically increased the stiffness of the trap by increasing the trapping power and observed that the maximum displacements traversed (**Extended Data Figs. 6b, 6c**) and the force traces for the bead within allY condensates do not show changes over 24 hours (**Extended Data Figs. 6f, 6g**). In contrast, the maximum distance traveled as well as the maximum force experienced by the particle increased for allY^20GtoS^ condensates over time (**Extended Data Figs. 6d, 6h, 6e, 6i**). This indicates that the restoring force responsible for the particle recoil in aged allY^20GtoS^ condensates is predominantly elastic. Taken together with the viscoelastic moduli estimated using MSD data (**Extended Data Figs. 5d, 5e, 5f)**, we conclude that physically aged condensates are terminally elastic solids. Note that aging was promoted by mutations to spacers, and hence the extent of crosslinking between different condensates should not change. However, the spacer-mediated compliance of the networks can change, and this enables facile transitions to terminally elastic materials.

### Terminally elastic condensates are non-fibrillar, semi-crystalline solids

Next, we asked if dynamical arrest induces changes to the mesoscale morphologies of condensates. Differential interference contrast (DIC) microscopy (**Fig. 5a**) shows that despite undergoing a transition from a Maxwell fluid to a Kelvin-Voigt solid (**Figs. 4c, 4d**), the micron-scale appearance of nascent versus aged condensates formed by allY^20GtoS^ remain essentially the same (**Fig. 5a**). Previous studies showed that fluid-to-solid transitions can be accompanied by the formation of amyloid fibrils ^30, 31^. To test for this possibility, we applied the amyloid-sensitive dye ThT to aged condensates formed by different sequence variants of A1-LCD. While this dye shows positive staining for fibrils formed by a positive control, insulin, at *t*_obs_ ≈ 24 hrs, it does not show positive staining for any of the aged PLCD condensates studied in this work (**Fig. 5b**). These results suggest that fluid-to-solid transitions observed for condensates formed by variants such as allY^20GtoS^ result in the formation of non-fibrillar, elastic solids.

**Fig. 5:**
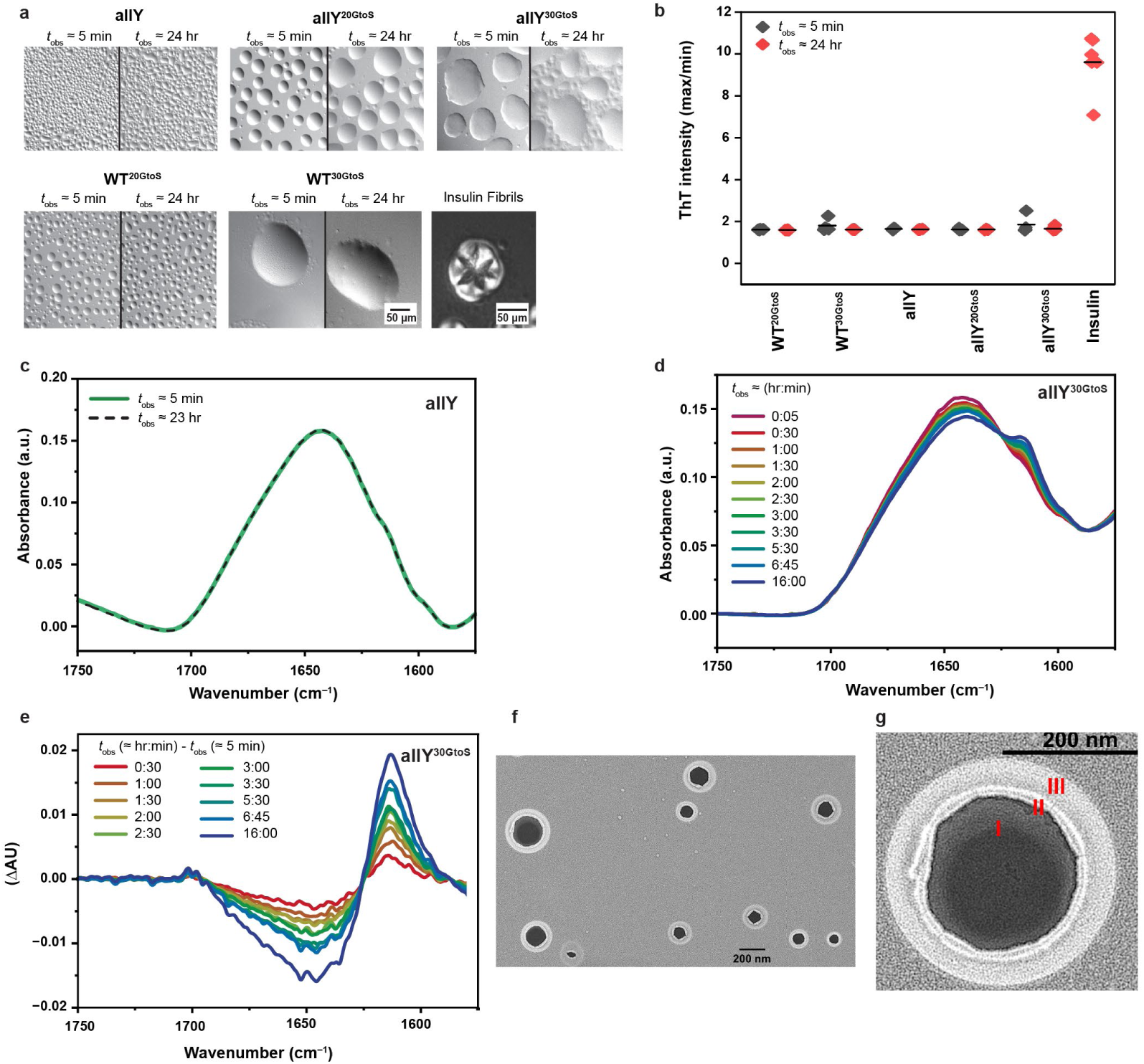
Disorder-to-order transitions drive the formation of beta-sheet-containing non-fibrillar, semi-crystalline solids as *t*_obs_ increases. (**a**) Representative images from DIC microscopy of different condensates at *t*_obs_ ≈ 5 min and at *t*_obs_ ≈ 24 hrs. Insulin fibrils are shown as positive control. **(b)** ThT fluorescence ratios of the maximal / minimal intensities for freshly prepared condensates and condensates at *t*_obs_ ≈ 24 hrs for various A1-LCD variants. No changes were observed in ThT staining for condensates formed as a function of aging. As a positive control, we show data for amyloid fibrils formed by insulin after 24 hrs from sample preparation. (**c)** FTIR spectra for allY condensates at *t*_obs_ ≈ 5 min and *t*_obs_ ≈ 23 hrs. (d) *t*_obs_-dependent FTIR spectra for condensates of allY^30GtoS^. **(e)** Difference spectra referenced to *t*_obs_ ≈ 5 min, show a decrease in signal for non-beta-sheeted regions and an increase in signal for beta-sheets, specifically aggregated beta-sheets (see **Extended Data Fig. 7a**). The difference spectra also reveal the presence of an isosbestic point, which is suggestive of a two-state disorder-to-order transition. **(f)** Freeze fracture deep-etch EM images of condensates of allY^30GtoS^ imaged at *t*_obs_ ≈ 24 hrs. **(g)** Zoom-in image of a semi-crystalline material formed by allY^30GtoS^. The semi-crystalline morphology is annotated by the presence of three regions labeled I, II, and III, which refer to an amorphous core (I), a lamellar crystalline region (II), and a halo generated by the reorientation of the samples during imaging, which originates due to the softness of the materials.

Next, we used Fourier transform infrared (FTIR) spectroscopy to study condensates at different values of *t*_obs_. Details of how the samples were prepared for these measurements are described in the Supplementary Material. Peaks in FTIR spectra, and the signals at different wave numbers can be referenced to extant basis sets to assign the contents of different types of secondary structures (**Extended Data Fig. 7a**). Condensates characterized at *t*_obs_ ≈ 5 min are disordered as evidenced by the equivalent contributions of different structural motifs to the measured FTIR spectra (**Extended Data Figs. 7b, 7c, 7d** and **7e**). While the FTIR spectra change minimally for allY over a 24-hour period (**Fig. 5c**), the spectra for allY^30GtoS^ show a clear change as a function of time (**Fig. 5d**).

Time-dependent changes in FTIR spectra for allY^30GtoS^ are consistent with a disorder-to-order transition. To emphasize this point, we plotted difference spectra, referenced to *t*_obs_ ≈ 5 min. We observed an increase in beta-sheet content and the changes are concordant with a two-state transition ^62^ as evidenced by the presence of an isosbestic point (**Fig. 5e**). Our positive control, insulin amyloid fibers that form upon overnight incubation, show FTIR spectra that bear qualitative resemblance to the aged condensates of allY^30GtoS^ (**Fig. 5d** versus **Extended Data Fig. 7h**). However, there are quantitative differences, and this is made clear by plotting the second derivative spectra, and the contributions of different components of the basis sets to the second derivatives. The area under the peak corresponding to “aggregated beta sheets” increases with *t*_obs_ for allY^30GtoS^ condensates (**Extended Data Figs. 7e, 7f, 7g**). However, the beta-sheet-contents of insulin amyloid fibers, which are known to feature cross-beta-sheet architectures (**Extended Data Figs. 7h, 7i**), is more than two-fold higher when compared to the condensates of allY^30GtoS^ that were studied at equivalent values of *t*_obs_. In general, the beta-sheet-contents increase with time, and this represents a disorder-to-order transition as a function of *t*_obs_ (**Extended Data Fig. 7j**). However, the DIC images and lack of ThT staining suggest that the supramolecular structures are not those of amyloid fibers that originate from cross-beta-sheeted networks. To gain further insights, we used freeze-fracture, deep-etch electron microscopy (EM) to probe the supramolecular organization of terminally elastic condensates formed by allY^30GtoS^ (**Fig. 5f**). The structures reveal a dense dark region that is consistent with an amorphous as opposed to crystalline organization (region I in **Fig. 5g**). However, the material is semi-crystalline, as evidenced by the lamellar ordering at the interface (region II in **Fig. 5g**). The materials are soft, and this causes a roll over that creates a halo as the sample is reoriented for deep etch EM imaging (region III in **Fig. 5g**). Overall, the picture that emerges is of beta-sheet-containing, non-fibrillar semi-crystalline materials that form as the condensates age. The disorder-to-order transitions and the semi-crystallinity lead to the terminally elastic, Kelvin-Voigt solids that we inferred from our passive and active microrheology measurements.

## Discussion

In this work, we have provided systematic assessments, via joint computational and experimental investigations, of sequence-specific and age-dependent viscoelasticity of condensates formed by PLCDs. We find that terminally viscous condensates are Maxwell fluids. The crossover timescale between dominantly elastic (short timescale) and dominantly viscous (long timescale) behaviors is governed by the strengths of sticker-mediated interactions. The storage and loss moduli increase by at least an order of magnitude as sticker strengths increase.

On a technical note, the LaSSI model used in this work employs a learning paradigm as opposed to a one-size-fits-all transferrable model. Specifically, we use a machine learning approach ^65^, which for single component systems leverages available experimental data such as small-angle X-ray scattering data ^35^ for the single chains in the one-phase regime ^8^. Alternatively, atomistic simulations of the *R*_g_ distributions with Tyr-to-Trp and Phe-to-Trp substitutions will suffice. However, at this juncture, neither input is available because of the ultra-low saturation concentrations associated with systems featuring multiple Trp residues. Molecular mechanics-based simulations of Trp containing systems are of questionable accuracy because of the double-ring systems, and the polarization effects induced by Trp. Absent a reliable parameterization of the interactions involving Trp residues, we cannot perform coarse-grained LaSSI simulations for variants where Phe and Tyr residues are replaced with Trp. Our preliminary simulations based on polarizable models suggest that Trp, as a sticker, has an effective valence that is greater than one^66^. This hypothesis, which explains why Trp, like other systems with five- and six-membered rings such as purine nucleobases ^67, 68^ make for stronger stickers than Tyr and Phe, will need systematic measurements of thermodynamic parameters and additional calibrations.

Both computations and measurements reveal a strong one-to-one correspondence between bulk viscosity within condensates and *c*_sat_, which quantifies the driving forces for phase separation. In computations, all the data collapse onto a single master curve without any rescaling or adjustments. The computations provide a purely thermodynamic description of condensates and the driving forces for condensation. The experimental data show a maintenance of the one-to-one correspondence, but instead of the data collapsing onto a single master curve, we find system-specific inverse correlations between viscosity and *c*_sat_. These deviations are attributable to hydrodynamic effects within condensates, and comparative analyses allow us to uncover the magnitudes of these effects. The physical aging of condensates can be accelerated by mutations to spacers, which also weaken the driving forces for forming terminally viscous condensates. Recent coarse-grained molecular dynamics simulations have established a one-to-one correspondence between the radii of gyration (*R*_g_) of disordered proteins in dilute phases and the viscosities in condensates ^63^. There is also a well-established positive, one-to-one correspondence between *c*_sat_ values and dilute phase *R*_g_ values ^8, 34, 35, 64^. Clearly, computations based on coarse-grained models predict a strong, master-curve-like one-to-one correspondence between viscosity and *c*_sat_. However, while the strong negative correlation is preserved in the experiments, the quantitative one-to-one correspondence is in fact system-specific. Simulations and theories that might be able to explain this observation are currently unavailable. Our findings provide the impetus for further exploring the origins of the system-specificity of hydrodynamic interactions.

Upon aging, we observe a disorder-to-order transition leading to semi-crystalline, non-fibrillar, beta-sheet-containing terminally elastic materials that are congruent with being Kelvin-Voigt solids. Our data are concordant with terminally viscous fluid-like states of condensates being metastable, disordered phases. Disorder-to-order transitions enable a conversion from metastable fluids to globally stable solids. Interestingly, in the absence of mutations to spacers, the activation barriers for network reconfiguration of fluid-like phases can be prohibitive, being on the order of 30 *RT* or twice as large depending on the sticker identities. This suggests that naturally occurring sticker and spacer grammars ensure enhanced metastability of PLCD condensates, which might explain why fluid phases are so readily observed in live cells, and why considerable attention has focused on the functional relevance of fluid forms of condensates.

Based on the totality of our observations, we propose that the condensation of PLCDs is best described using a bistable model whereby the free energies of dense phases are written in terms of an order parameter (Φ | *c*_dense_), which quantifies the extent of structural ordering on the manifold of fixed density (*c*_dense_) (**Fig. 6**). We propose that terminally viscous Maxwell fluids are metastable, disordered phases defined by low values of the order parameter. Fluid-to-solid transitions are disorder-to-order transitions and the likelihood of realizing this transition will be governed by the height of the barrier between the Maxwell fluid and Kelvin-Voigt solid (**Fig. 6**). Our current data are consistent with the metastability of fluid states being weakened by mutations to spacers.

**Fig. 6:**
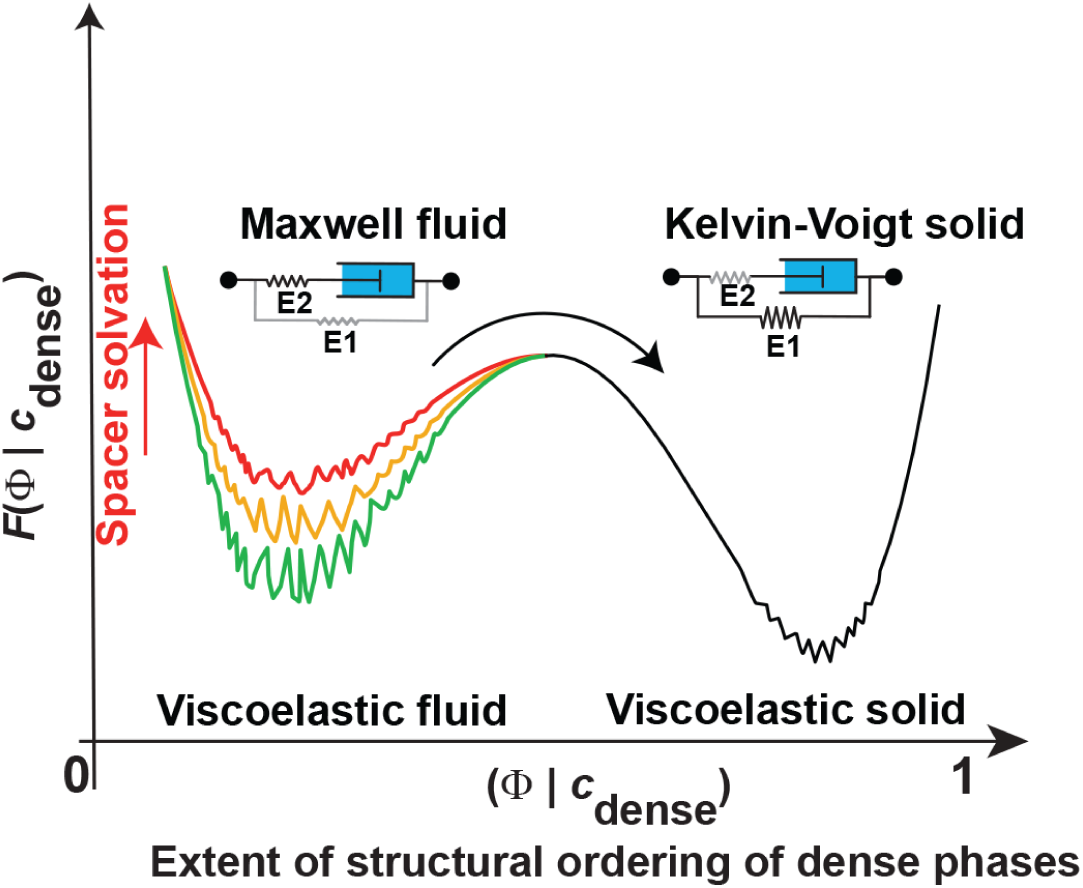
Proposed free energy profile showing bistability and intra-well ruggedness. The well corresponding to low values of the order parameter is a disordered, terminally viscous Maxwell fluid. The well on the right, corresponding to high values of the order parameter is the ordered, Kelvin-Voigt solid. The barrier for disorder-to-order transitions is expected to be high, > 30*RT*, for condensates formed by naturally occurring PLCDs. This barrier is lowered, by destabilizing fluid-like states via mutations to spacers that increase their effective solvation volumes. Within the metastable and globally stable wells, we expect there to be ruggedness due to polymorphisms in the solid phase, and the contributions from conformational heterogeneity in the fluid phase.

The free energy profiles in **Fig. 6** are annotated by two features. The terminally fluid and solid phases are represented using a standard linear solid or Zener model ^69^. In this model, a time-dependent rise of the elasticity of the E1 element captures the transitions from a dominant Maxwell-type behavior of nascent condensates (E1 < E_2_) to a Kelvin-Voigt-type behavior as condensates age (E1 >> E_2_). We also allow for the possibility that increased metastability of the fluid state arises from ruggedness within the well that corresponds to low values of Φ. Our proposal represents a combination of the Landau picture ^70^ of transitions along (Φ | *c*_dense_) and the Zwanzig picture ^71^ of roughness within a well.

Our findings and the resultant proposals stand in contrast to those of Jawerth et al. They used active microrheology and fluorescence recovery after photobleaching to characterize condensates formed by the protein PGL3 ^12^. Jawerth et al., found that fluid-like condensates formed by this protein are Maxwell fluids, a result we recapitulate for PLCDs. They also found that aged condensates continue to behave as Maxwell fluids, even though the condensates no longer fused, and rheology measurements suggested that the condensates were dynamically arrested upon aging. Based on their observations, Jawerth et al., proposed that the PGL3 condensates were “Maxwell glasses” defined as Maxwell fluids showing age-dependent terminal relaxation times ^12^. Our data, based on direct measurements of terminal states, leads to a different picture. We propose that viscoelastic Maxwell fluids are metastable. Weakening the metastability of terminally viscous states unmasks the transition to a globally stable, terminally elastic state, which for the PLCD systems appears to be a Kelvin-Voigt, non-fibrillar solid.

Recent bioinformatics analyses showed an inverse correlation between sticker and spacer identities ^35^. Increasing the number of Tyr stickers is positively correlated with increased numbers of Gly residues as spacers. In contrast, increased numbers of Phe stickers are accompanied by increased number of Ser residues as spacers. Measurements of PLCDs that form solids in cellular systems have also documented a preference for high numbers of Phe stickers and Ser residues as spacers ^72^. Taken together, these trends suggest an evolutionarily encoded preference whereby there is either a selection for enhanced metastability of fluid-like phases (positive correlation between Tyr and Gly contents) or a selection for facile transitions to solid phases (positive correlation between Phe and Ser contents). How this plays out for condensates formed by mixtures of PLCDs with compositionally distinct preferences will need further investigation ^9^. Importantly, being able to compute moduli from simulations of the network structures of condensates ^8^ should allow us to design PLCD sequences with bespoke material properties ^9, 73^.

Previous work showed that the linear patterning of aromatic stickers affects the formation of fluid-versus solid-like PLCD condensates ^34, 74^. In full-length proteins with PLCDs, Wang et al. showed that mutations to spacers can have a direct impact on the time-dependent changes to mobilities in dense phases^21^. Given the large barriers we estimate for network rearrangements (**Fig. 2c, 2d**), it appears that for many naturally occurring systems, including PLCDs, the barrier to conversion from terminally viscous fluids to terminally elastic solids might be insurmountable on biologically relevant timescales, especially for those that are turned over rapidly. This might explain why “dynamicity” is often invoked as a key determinant of condensate functions in cells ^16, 18, 19^. In contrast to condensates that turn over, there are long-lived condensates such as Balbiani bodies ^75^, where the sticker grammars and choice of spacers appear to enable facile conversion to terminally elastic solid phases ^74^. Likewise, the sequence grammars of scaffolds of solid phase condensates such as A-bodies in nuclei of stressed cells ^76^ might be such that the barriers to conversion from fluids to solids is readily negotiated on functionally relevant timescales. Our work highlights the need to uncover the connections between sequence-encoded interactions and the time-dependent material properties of condensate-forming systems both in vitro and in vivo. Our work also invites detailed structural characterizations of metastable fluids and globally stable solid phases, to investigate how the interplay of homotypic and heterotypic interactions influence material states and structures in multicomponent mixtures ^9, 13, 23, 77^.

## Acknowledgments

This work was supported by the US National Institutes of Health through grants R01NS121114 (T.M. and R.V.P), R35 GM138186 (P.R.B), and the St. Jude Children’s Research Collaborative on the Biology and Biophysics of RNP Granules (P.R.B., T.M., and R.V.P). S.R.C. acknowledges support from the National Institutes of Health (T32 EB028092). We thank George Campbell from the Cell and Tissue Imaging Center at SJCRH, which is supported by SJCRH and NCI (grant P30 CA021765) for assistance with the DIC and confocal microscopy. We acknowledge the Washington University Center for Cellular Imaging (WUCCI), which is supported by the Washington University School of Medicine, The Children’s Discovery Institute of University and St. Louis Children’s Hospital (CDI-CORE-2015-505 and CDI-CORE-2019-813) and the Foundation for Barnes-Jewish Hospital (3770 and 4642).

## Author contributions

Conceptualization: P.R.B., R.V.P., and T.M.; Methodology: P.R.B., I.A., W.M.B., S.R.C., R.V.P., and T.M.; Investigation: I.A., W.M.B., S.R.C., A.E.P., M.F., A.S., A.B., G.S., D.T.T., P.R.B., R.V.P., and T.M.; Resources: P.R.B., R.V.P., and T.M.; Writing – original draft and revisions: P.R.B., R.V.P., and T.M.; Writing – reviewing and editing: all authors; Funding acquisition: P.R.B., R.V.P., and T.M.

## Data and materials availability

All data are available in the manuscript or the supplementary materials. All expression plasmids are available from T.M. under a material transfer agreement with St. Jude Children’s Hospital. Codes for microrheology data analysis are available on GitHub (see https://github.com/BanerjeeLab-repertoire/Material-properties). All simulation results and the code for the computation of moduli are available via the Pappu Lab Github (see https://github.com/Pappulab/material-properties).

---

**Extended Data Fig. 1.**
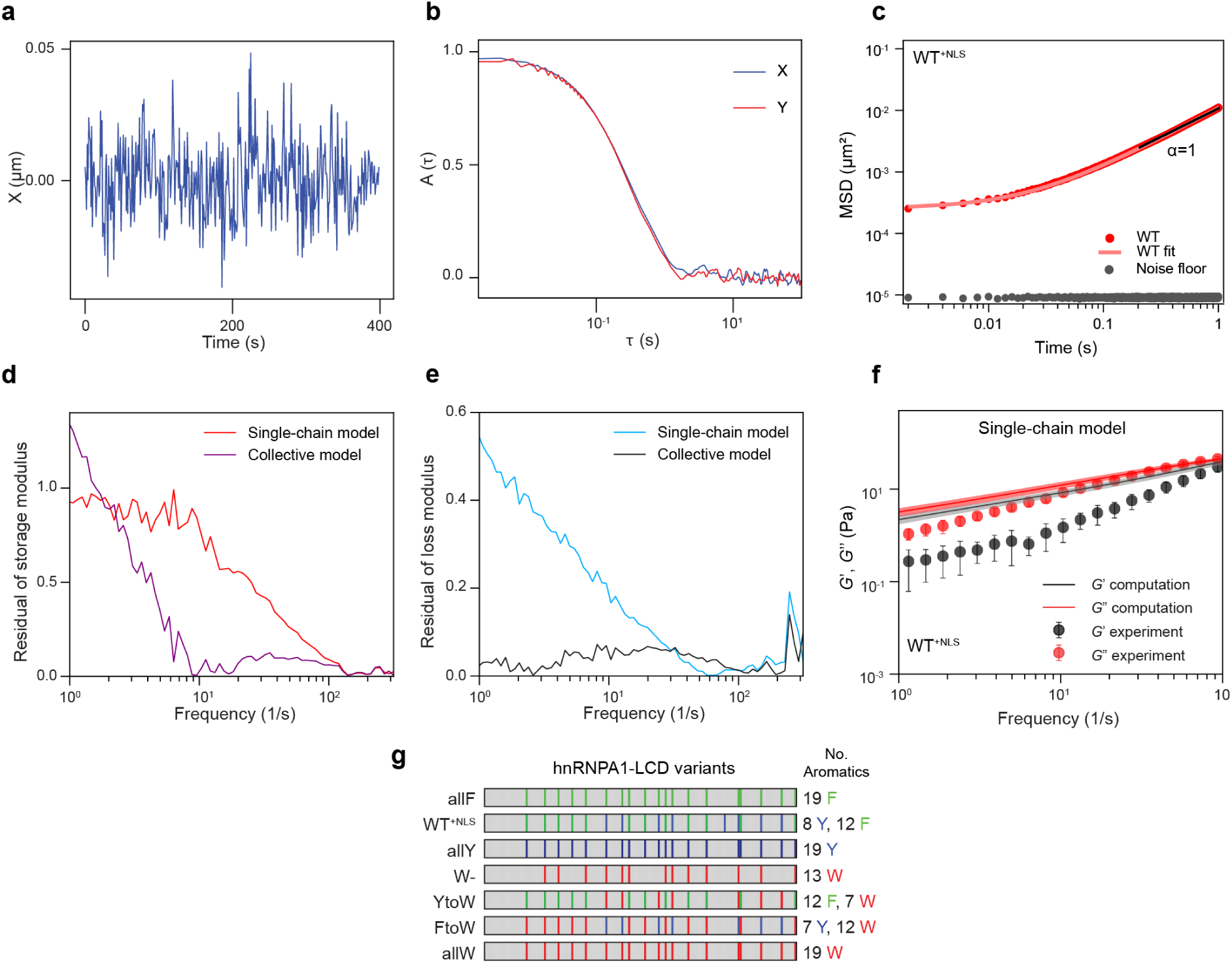
**(a)** Representative positional trajectory, tracked along the horizontal axis, of a bead thermally fluctuating within the optical trap inside the condensate. **(b)** The normalized autocorrelation function of positional fluctuations of a bead, confined by the optical trap, inside a condensate. The correlation functions measured along the horizontal and vertical axes are shown. **(c)** Ensemble-averaged MSD of 200 nm beads in condensates formed by WT^+NLS^. The solid line is a fit to the equation *MSD*(*τ*) = 4*Dτ^α^* + *N* and reveals that at longer timescales (>10ms) the beads are diffusive. However, at shorter timescales (<10 ms), the beads are sub-diffusive, indicating that the condensates are dominated by an elastic response. **(d)** Residuals between computed and measured storage moduli for WT^+NLS^ condensates. These residuals are shown for the single-chain model (red) and collective model (magenta). The residuals were computed as the absolute values of differences between the logarithms of the computed and measured moduli. Values of zero imply exact matches, values of one imply a discrepancy of an order of magnitude, and so on. At low frequencies, the sharp increase in residuals for the collective model is due to increased noise in the experimentally derived storage moduli. **(e)** Residuals between computed and measured loss moduli using the single chain (blue) or collective chain models (black). The residuals were computed as in panel (d). **(f)** Comparison of computed and measured dynamical moduli for WT^+NLS^ condensates. The computations use the single-chain model. The computed crossover frequency was rescaled to match the experimental value. No other adjustments or fitting of the data was performed. The inadequacy of the single-chain model is made clear by the fact that this model overestimates the storage modulus by almost two orders of magnitude. Similar overestimates were found for other systems, and they stand in contrast to the results obtained using the collective model shown in Fig. 1b. **(g)** A schematic diagram of the constructs deployed in the various measurements in this work is summarized. Grey represents non-aromatic spacer residues, whereas colored stripes represent the locations of the indicated aromatic residues.

**Extended Data Fig. 2.**
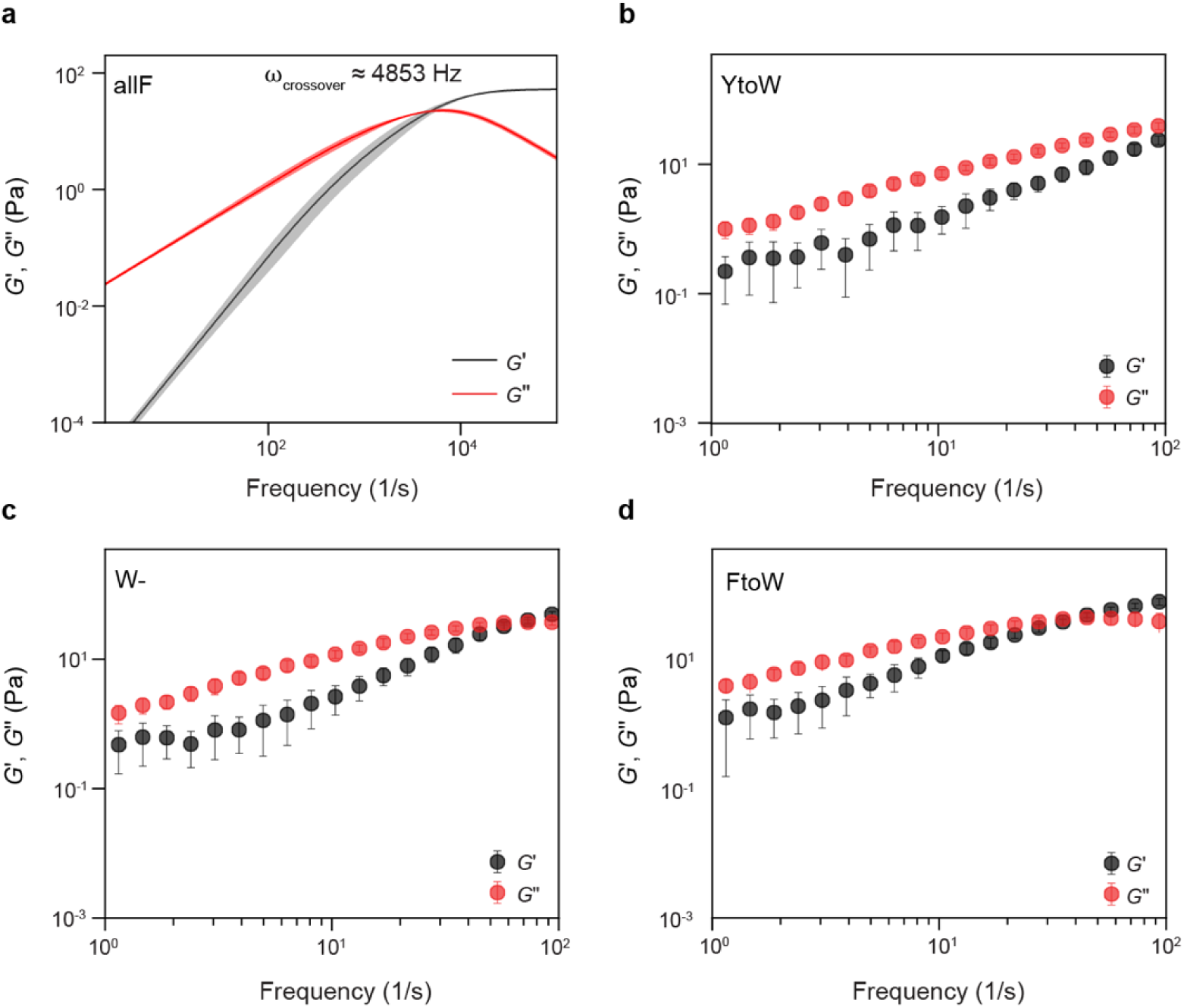
**(a)** Computed dynamical moduli for condensates formed by allF. The crossover frequency analysis of the simulation results was performed using the collective model. The crossover frequency was calculated numerically by estimating where the curves corresponding to the storage and loss modulus intersect. The value of ω_crossover_ obtained by extrapolation is indicated in the legend. The error band for the computed data represents the standard deviation from 3 replicates. **(b)** Measured dynamical moduli (symbols) for condensates formed by the YtoW variant. Error bars represent the standard deviation as calculated from the moduli of 20 condensates. **(c, d)** Measured dynamical moduli for condensates formed by W- and FtoW variants, respectively. Error bars are standard deviations as calculated from the moduli of 20 condensates.

**Extended Data Fig. 3.**
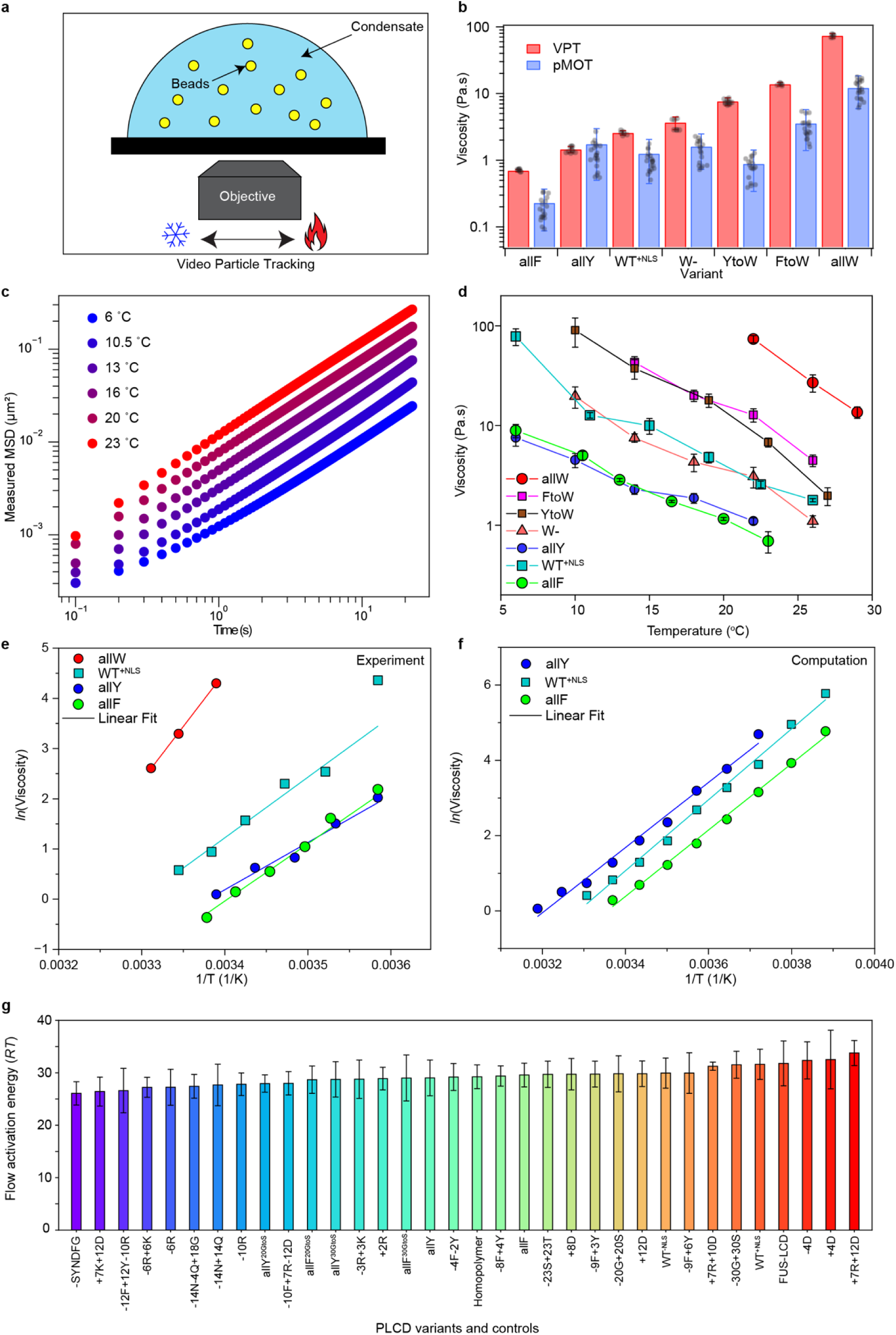
**(a)** Schematic for the temperature-dependent VPT measurements. Diffusion of multiple 200 nm polystyrene beads is recorded at distinct temperatures. **(b)** Room-temperature bulk dynamical viscosity of condensates formed by different sequence variants as measured by VPT compared to pMOT. Data were collected from at least five trials featuring at least ∼50 tracked particles per trial. Error bars represent ±1 standard deviation. Although the viscosity values obtained from these two independent experimental measurements are in good agreement with one another, we note that the terminal zero-shear viscosity, obtained from the slope of the frequency-dependent loss modulus in pMOT measurements, is an asymptotic property, which means that this quantity is difficult to estimate for viscoelastic materials when the elastic and viscous moduli become similar, even below the crossover frequency. The low-frequency limit of the measurement (about 0.1 Hz which corresponds to ∼10s) may not be low enough to estimate the zero-shear viscosity, especially for condensates with high viscoelasticity, such as for variants with Trp residues as stickers. This explains the minor discrepancy in the viscosity values obtained from VPT and pMOT measurements. **(c)** Representative plots of the ensemble-averaged MSD of 200 nm polystyrene beads within condensates formed by allF. Measurements were performed using VPT-based microrheology at different temperatures as set by a custom-made thermal stage attached to the microscope. **(d)** Measured temperature-dependent viscosities for the condensates formed by different sequence variants of A1-LCD. **(e)** Arrhenius plots of measured viscosities for condensates formed by allW, WT^+NLS^, allY, and allF. **(f)** Arrhenius plots using temperature-dependent viscosities from simulations analyzed using the collective model. Data are shown for condensates of allF, WT^+NLS^, and allY. **(g)** Flow activation energies estimated from computations. These energies are shown in units of *RT*, where *T* = 25 °C and *R* is the ideal gas constant in units of kcal / mol-K. The variant names were taken from the work of Bremer et al., (ref. 35) and Farag et al., (ref. 8).

**Extended Data Fig. 4.**
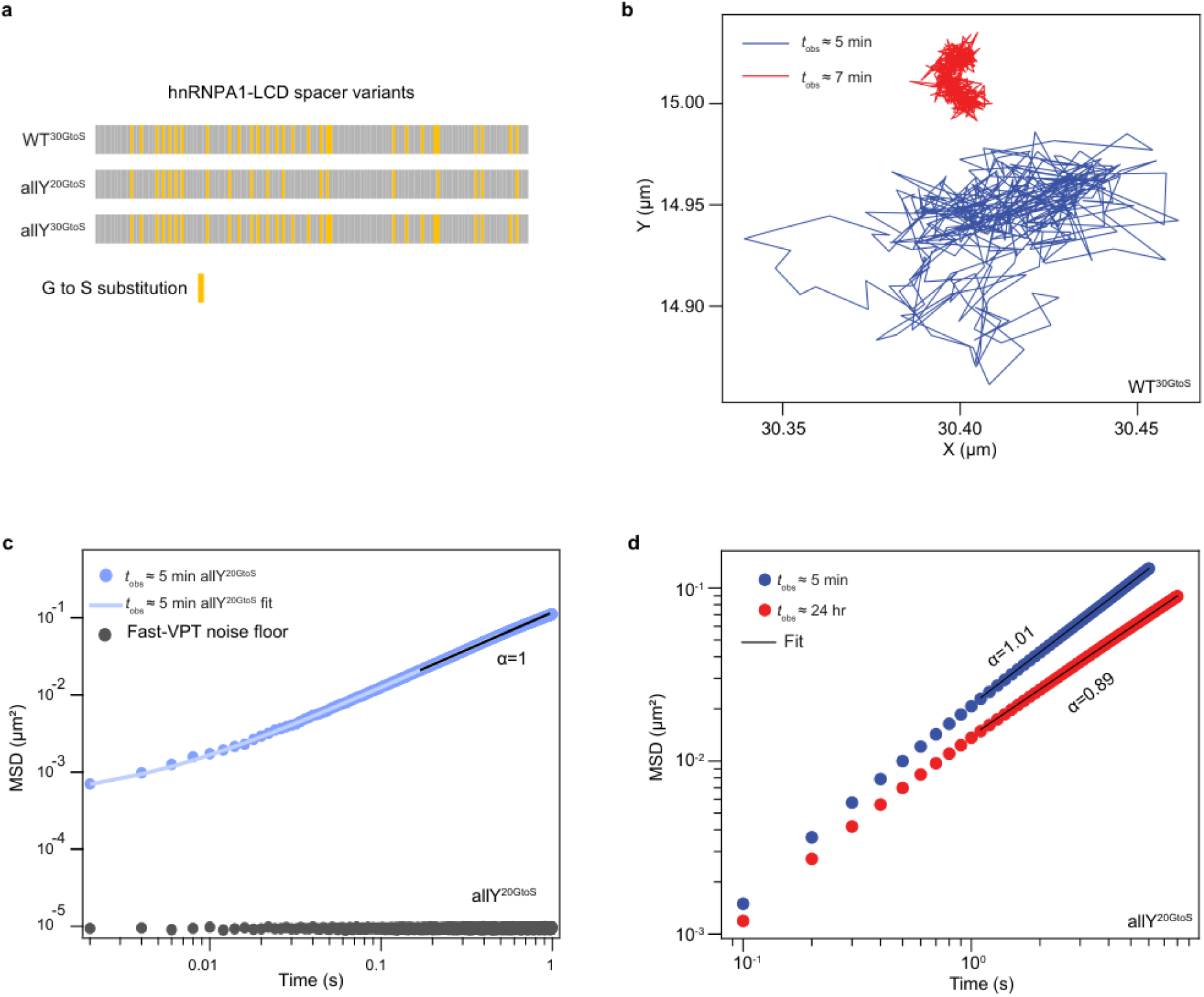
**(a)** Schematic denoting the position of Glycine to Serine substitution in the WT^+NLS^ and allY sequence backgrounds. **(b)** Trajectories of a 200 nm bead within condensates of WT^30GtoS^. Two different trajectories are shown, one for *t*_obs_ ≈ 5 min (blue) and the other for *t*_obs_ ≈ 7 min (red). These data show that WT^30GtoS^ condensates undergo rapid dynamical arrest within a short time. (**c**) Ensemble-averaged MSD of 200 nm beads diffusing within allY^20GtoS^ condensates tracked at a high acquisition rate of 500 fps. The solid blue line is a fit to the equation *MSD*(*τ*) = 4*Dτ^α^* + *N*. Also plotted is the average noise floor (black). These data show that allY^20GtoS^ condensates are viscoelastic Maxwell fluids at small *t_obs_*, with a terminal behavior of a viscous fluid (α = 1). **d)** Ensemble-averaged MSD of 200 nm beads (tracked at an acquisition rate of 10 fps) in condensates formed by allY^20GtoS^. The MSDs were measured at different values of *t*_obs_. Solid lines are the fits to the equation *MSD*(*τ*) = 4*Dτ^α^* + *N*. Also indicated are the values of the exponents obtained from the fits, which show a clear deviation from the linear behavior for the aged condensates, and the onset of sub-diffusive motions.

**Extended Data Fig. 5.**
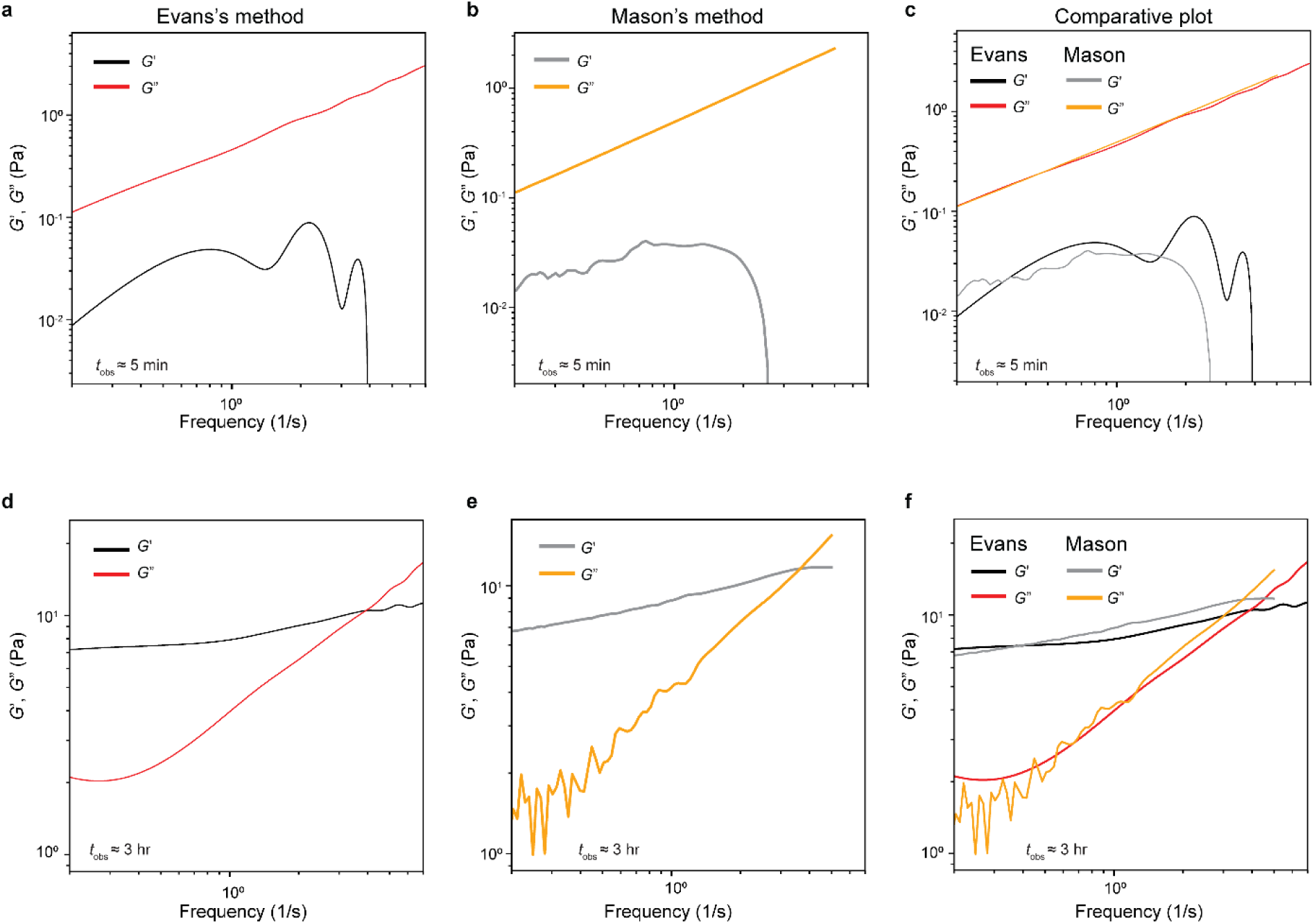
**(a-c)** Storage (*G*′) and loss (*G*″) moduli of allY^30GtoS^ condensates at *t*_obs_ ≈ 5 min as calculated from the measured MSDs of embedded 200 nm probe particles (Fig. 3f in the main text). The calculations were performed following the Evans method **(a)** and the Mason method (**b**, see Supplementary Material). A comparison between the results obtained from the two methods is shown in **(c)**. **(d-f)** Same as **(a-c)** but for allY^30GtoS^ condensates at *t*_obs_ ≈ 3 hrs.

**Extended Data Fig. 6.**
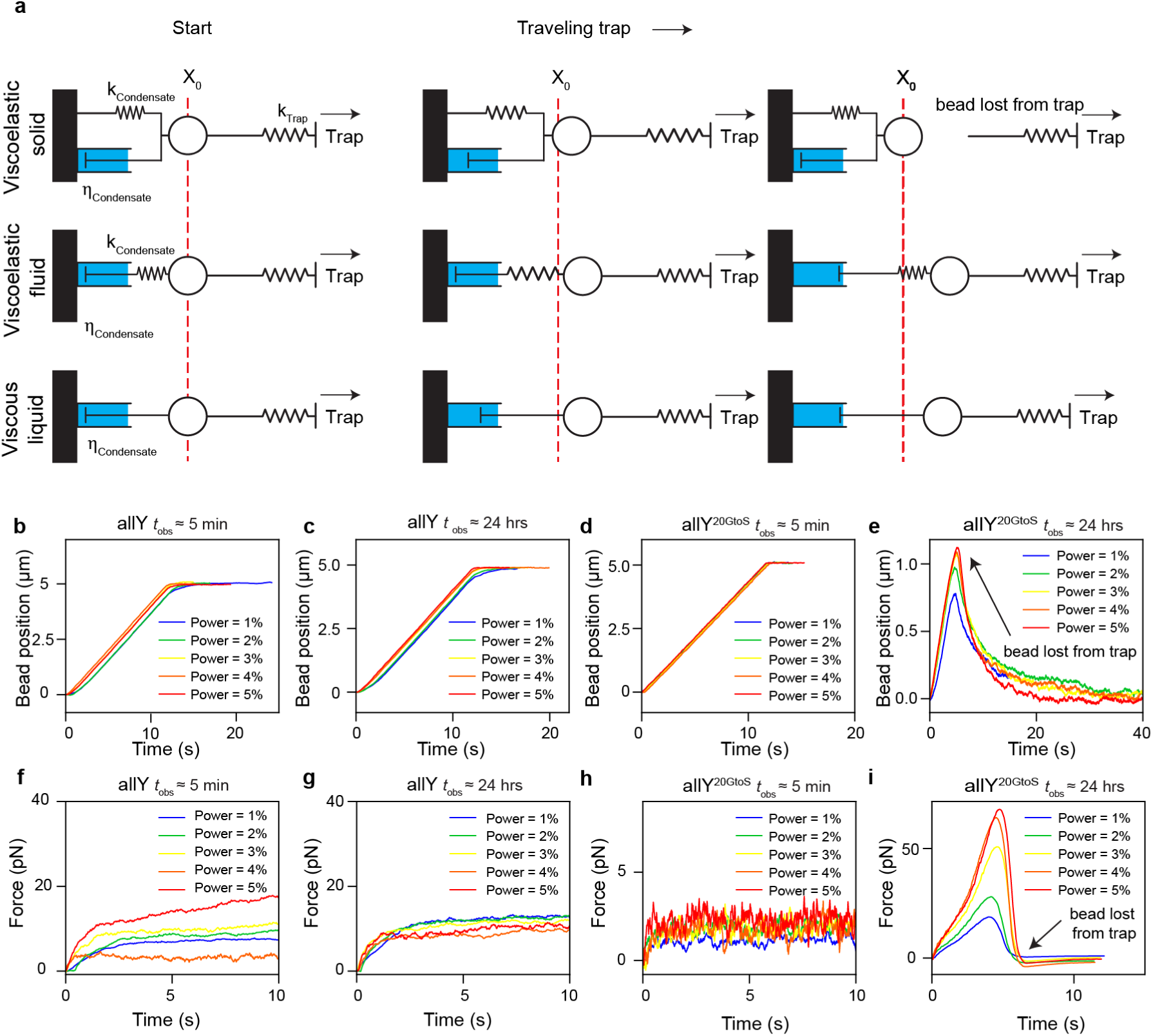
**(a)** Schematic showing the expected outcomes of a particle embedded within a viscoelastic Kelvin-Voigt solid (top), viscoelastic Maxwell fluid (middle), and viscous liquid (bottom) upon creep deformation by an optical trap. As the optical trap exerts a force to drag a particle embedded within the material, the dashpot element applies a dissipative drag force while the spring element applies a restoring force. Except for the viscoelastic solid, the creep deformation is irreversible (the final bead position is distinct from the initial bead position X_0_, indicated by the red vertical line). In the case of a viscoelastic solid, the bead recoils back to its initial position due to the stored energy within the elastic element. In both viscoelastic fluids and viscous liquids, the deformation energy is eventually dissipated through the dashpot or the viscous element, giving rise to a terminally viscous behavior. **(b)** Bead position as a function of time in response to the programmed trap movement in allY condensates at *t*_obs_ ≈ 5 min at increasing power levels of the trapping laser. **(c)** Same as (**b**) but for the allY condensates at *t*_obs_ ≈ 24 hrs. **(d)** Bead position as a function of time in response to the programmed trap movement in allY^20GtoS^ condensates *t*_obs_ ≈ 5 min at increasing power levels of the trapping laser. **(e)** Same as (**d**) but for the allY^20GtoS^ condensates at *t*_obs_ ≈ 24 hrs. **(f)** The force exerted on the bead by the allY condensate network in (**b**). **(g)** The force exerted on the bead by the allY condensate network in (**c**). **(h)** The force exerted on the bead by the allY^20GtoS^ condensate network in (**d**). **(i)** The force exerted on the bead by the allY^20GtoS^ condensate network in (**e**).

**Extended Data Fig. 7.**
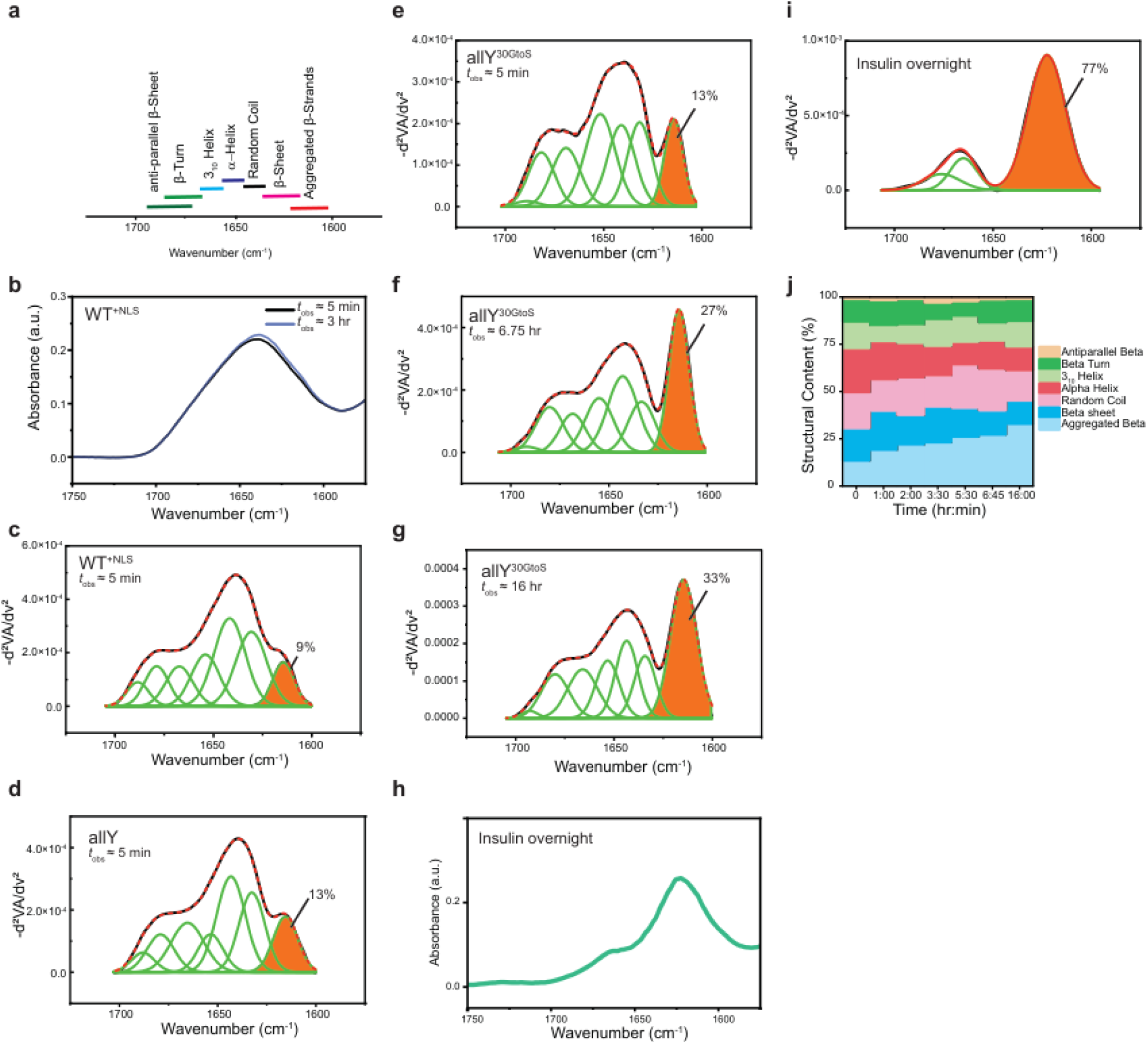
**(a)** Peaks in FTIR spectra correspond to the presence of specific types of secondary structures. **(b)** FTIR spectra for condensates formed by WT^+NLS^. The spectra are shown for *t*_obs_ ≈ 5 min and *t*_obs_ ≈ 3 hrs. (**c**) Inverse second derivatives of the measured FTIR spectra and contributions from different members of the basis set. The orange shading quantifies the area under the peak that corresponds to aggregated beta sheets. Data are shown here for WT^+NLS^ condensates for *t*_obs_ ≈ 5 min. (**d**) The inverse second derivative of FTIR spectra and contributions from different members of the basis set to data obtained for allY condensates at *t*_obs_ ≈ 5 min. (**e**) – (**g**) Second derivative of FTIR spectra obtained at different values of *t*_obs_ for condensates formed by allY^30GtoS^. (**h**) FTIR spectra for amyloid fibers of insulin collected at *t*_obs_ ≈ 24 hrs. (**i**) Second derivative of the measured FTIR spectra for insulin fibers and contributions from different members of the basis set. The orange shading quantifies the area under the peak that corresponds to aggregated beta sheets. **(j)** Stacked plots of secondary structure contents for allY^30GtoS^ condensates plotted as a function of time, where the abscissa refers to different values of *t*_obs_.

