## Supplementary Methods for "Sequence-specific interactions determine viscoelasticity and aging dynamics of protein condensates"

### Materials and Methods

#### Protein construct design and purification

All A1-LCD constructs were based on residues 186-320 of human hnRNPA1 (UniProt: P09651; Isoform A1-A). The coding sequences for the variants were synthesized (by ThermoFisher) with the inclusion of the coding sequence for an N-terminal TEV cleavage site and flanking attB sites. The sequences were then subcloned into a pDEST17 vector using LR clonase (ThermoFisher). Proteins were expressed and purified (Supplementary Fig. S1) as previously described [1] with a few modifications. The purification of tryptophan-containing variants was modified as follows: the Nickel-affinity column buffers contained 6 M Urea instead of 4 M, TEV cleavage was conducted at room temperature with 3 M Urea instead of 2 M, and the GdmHCl concentration in the size exclusion buffer was increased to 4 M. Additionally, the protein concentration of the allW variant had to be kept below 50 μM during TEV cleavage to prevent precipitation. The amino acid sequences of each of the constructs are shown in **Table S1**.

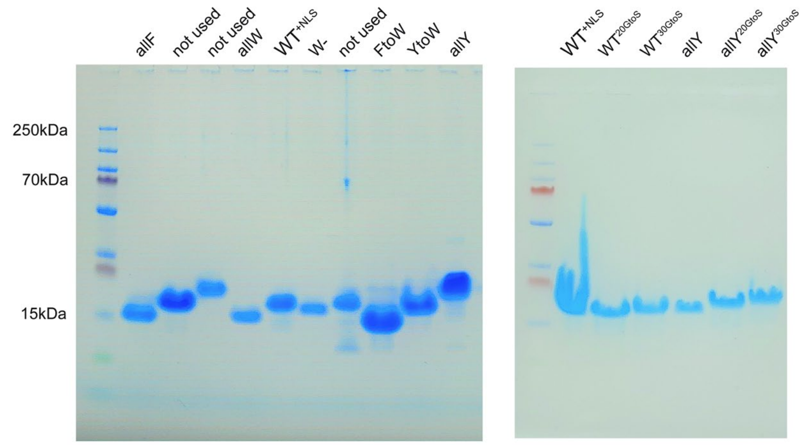


**Supplementary Figure S1.** SDS-PAGE (sodium dodecyl sulfate–polyacrylamide gel electrophoresis) gel image of purified proteins used for the experiments.

### Measurements to construct phase boundaries

Buffer exchange into native buffer without excess salt and measurements of phase boundaries were performed as described previously [2]. Phase separation of A1-LCD solutions was induced by adding NaCl to a final concentration of 150 mM. Dilute phase concentrations as a function of temperature were determined in triplicate as outlined in previous work [2, 3]. Briefly, the dilute phase was separated from the dense phase via centrifugation, diluted if the desired temperature was above ambient values, and the concentration was determined via standard UV-VIS absorbance measurements. Dilute phase concentrations at or above 30°C were determined via cloud point measurements using static light scattering as previously described [2, 4].

### Glass slide and coverslip preparation

Glass slides and coverslips were either coated with Tween20 or with Sigmacote (Sigma-Aldrich). To coat with Tween20, we followed the procedure of Alshareedah et al. [5] Briefly, glass slides and coverslips were immersed in a 20% vol/vol solution of Tween20 for 30 minutes. Next, the glass slides and coverslips were washed with milliQ water several times and subsequently dried with compressed air. For Sigmacote coating, the slides and coverslips were first cleaned with 70% ethanol and then coated with Sigmacote following the manufacturer's procedure.

### Sample preparation and chamber assembly

Samples were prepared by mixing appropriate volumes of 20 mM HEPES buffer pH 7 (adjusted with ammonium hydroxide), buffer exchanged protein, and yellow-green carboxylate-modified polystyrene beads (FluoSpheres, Invitrogen). The bead diameter was 0.2 µm, 1 µm, and 2 µm for video particle tracking, passive microrheology with optical tweezers, and creep test measurements, respectively. 1.5 M NaCl was then added to a final concentration of 150 mM to induce phase separation. 2 to 5 µL of the prepared sample was spotted onto a coverslip and sandwiched with a glass slide using 3 to 4 layers of double-sided tape. Mineral oil was then injected into the chamber to prevent the sample from evaporation by filling the remaining volume and isolating the sample from air.

### Temperature-controlled video particle tracking (VPT)

We prepared the protein condensates in a buffer that contained 200 nm carboxylate-modified yellow-green beads (Invitrogen) as described in the sample preparation section above. The sample was allowed to equilibrate for a few minutes before loading it onto the microscope for imaging. Fluorescence imaging of the sample was performed using a Zeiss Primovert inverted microscope equipped with a 40× air objective and a Teledyne FLIR blackfly S USB3 CMOS camera. The microscope also contained a custom-designed thermal stage (INSTEC) with a heat insert to fix and measure the temperature of the sample through a thermocouple. The exposure time was set to 100 ms, which is also the frame time of the movies (frame rate is 10 frames per second). Movies of the beads diffusing within the condensate were recorded for 2 to 5 minutes and saved for further analysis. For the room temperature viscosity measurements, all analyses were performed using at least six individual condensates through multiple sample preparations. For the condensate aging experiments, we used a 100× oil immersion objective. For the flow activation energy measurements, condensates were first equilibrated to the lowest experimental temperature for ~10 minutes. Movies of the diffusion of beads were then collected. The temperature was then increased to the next higher temperature and the sample was equilibrated at that temperature for ~10 minutes. This process was repeated until the upper temperature limit was reached, which was checked to be significantly lower than the upper cloud point temperature of the variants. In some cases, droplets fused on the coverslip and formed a continuous condensed phase that is a few micrometers in height. In such cases, we imaged several fields of view and analyzed them individually to extract viscosity. The viscosity values were averaged and the error in the mean values was estimated as the standard deviation. For the high-speed VPT experiments (**Extended Data Figs. 1d** and **4c**), the sample was prepared in an identical way to all other VPT samples and loaded on a Zeiss Axio Observer inverted microscope equipped with a 100× oil immersion objective. High speed imaging (200-500 frames per second) was achieved using the back-illuminated Kinetix 22 sCMOS camera (Teledyne Photometrics).

**Passive Microrheology with Optical Tweezers (pMOT)**

All pMOT experiments were conducted as described in detail by Alshareedah et al. [5] The theoretical basis of the method relies on the works of Preece et al., [6] Yao et al., [7] Tassieri et al., [8], as well as Mason and Weitz [9]. pMOT experiments were conducted on a LUMICKS C-Trap microscope equipped with a 60× water immersion objective. Condensates were prepared in a buffer containing 1.0 μm beads. Condensates were allowed to settle on the surface of the coverslip for ~20 minutes or until no fusion events were observed and no condensates were observed in the solution. Individual beads within condensates were trapped using the optical tweezer at minimum power (<100 μW). The trapped bead was positioned at the center of the condensate in the *xy* plane and ~3 μm above the glass surface. The bead trajectory was tracked using a built-in tracking algorithm through the bright field camera of the microscope at a rate of 500 Hz. Bead trajectories were collected for 5 to 20 minutes and saved for further analysis.

**Optical tweezer-based creep test**

The laser tweezer-based creep test was performed in a similar manner as pMOT experiments. However, instead of tracking the trajectory of a stationary bead, we programmed the optical tweezer to travel 5.0 μm at a speed of 0.5 μm/s. The optically trapped bead (2 μm in diameter) was imaged as it responded to the traveling trap and the bead trajectory within the field of view was tracked using the bright field camera. For each system, we performed the creep test at five different trapping laser powers (1−5 % with an increment of 1%, 10−50 μW). Prior to each creep test measurement, the trajectory of a stationary bead was collected for 2 minutes at the respective tracking power. This trajectory was used for the optical trap calibration (see *data analysis* section). The force experienced by the bead was obtained by measuring the displacement of the bead from the center of the optical trap using a quadrant photodiode.

### Data analysis

*Passive microrheology with optical tweezers and the creep test*

Trajectories of trapped beads were analyzed in two steps: (a) calibration of the optical trap, and (b) computation of the complex shear modulus also referred to as the viscoelastic modulus [5]. For calibration, trajectories in $x$ and $y$ were corrected for drift using a cubic spline-fitting algorithm to remove the long-time drift of the particle. Next, the variance of the positional fluctuations in the $x$ and $y$ directions was computed. The trap stiffness was then computed using the equipartition theorem [10]:

$$\kappa_x = \frac{k_B T}{\langle x^2 \rangle} \qquad (1)$$
$$\kappa_y = \frac{k_B T}{\langle y^2 \rangle}$$

where $k_B$ is the Boltzmann constant and $T$ is the temperature in Kelvin.

To calculate the complex shear modulus, we first computed the normalized position autocorrelation function $A(\tau)$. This was done via the multipletau Python library. Next, the autocorrelation function was Fourier transformed into the frequency domain $A(\omega)$ [8]. We next calculated the complex shear modulus using the relation:

$$G^*(\omega) = \frac{\kappa}{6\pi a}\left(\frac{i\omega A(\omega)}{1 - i\omega A(\omega)}\right) \qquad (2)$$

In (2), $\kappa$ is the trap stiffness in $x$ or $y$ dimension and $a$ is the trapped particle radius. After the analysis of data from all tested condensates (~20 condensates and ~40 one-dimensional trajectories), the values of $G'$ and $G''$ were averaged at each frequency and the error was estimated as the standard deviation. The top and bottom 5% of the data were excluded as outliers. More details on this method can be found in our earlier work [5].

For the creep test, the optical trap was calibrated using the equipartition theorem as described above by monitoring the trajectory of a bead in a stationary optical trap. Then, forces were computed by multiplying the trap stiffness by the position of the bead from the center of the optical trap, which is measured using a quadrant photodiode (LUMICKS, C-trap). The absolute bead position within the field of view was measured using a camera, which is then normalized by the bead coordinates at the start of the experiment before the trap motion is initiated. Both bead positions and forces were calculated and plotted using custom python codes (see *Software* section below).

*Temperature-controlled video particle tracking*

Video particle tracking (VPT) measurements monitor the movement of beads inside the condensates and output movies which can be further analyzed to get the mean square displacement (MSD) of the beads [11]. MSDs can be used to estimate other quantities of interest such as diffusion coefficients, diffusivity exponents, and viscosity. We first used the TrackMate plugin of ImageJ FIJI software to track the diffusion of beads and produce two-dimensional trajectories of each bead within the condensates [12]. Next, we calculated the diffusion of the center of mass of beads by computing the center of mass vector $\boldsymbol{R}$ at each time point. Calculation of the center of mass vector was performed using particle velocities according to the following equation:

$$\boldsymbol{R_{COM}}(k) = \boldsymbol{R_0} + \sum_{k=0}^{k} \boldsymbol{\bar{v}_k} = \boldsymbol{R_0} + \sum_{k=0}^{k} \left( \frac{1}{N_k} \sum_{i=1}^{N} \boldsymbol{v_{i,k}} \right) \Delta k \tag{3}$$

Here, $\boldsymbol{R_0}$ is the initial center of mass vector calculated by averaging the coordinates of all beads in the first frame ($k$=1), $\boldsymbol{\bar{v}_k}$ is the mean velocity of all particles in frame $k$, $\boldsymbol{v_{i,k}}$ is the velocity of particle $i$ in frame $k$ in units of μm/frame and $N_k$ is the number of particles in frame $k$. The quantity $\Delta k$ is the frame difference which is included in the equation to have correct units but is set to 1 since we are analyzing consecutive frames. This method of center-of-mass (COM) calculation from particle velocity is insensitive to the appearance or disappearance of particles from the field of view, which can lead to false COM shifts if calculated using the conventional definition. Once the COM is calculated, we subtract it from the individual bead trajectories to ensure the absence of any sample drift in the data. Following that, we calculate the ensemble and time averaged mean squared displacement (MSD) using:

$$MSD(\tau) = \langle \boldsymbol{R}(t+\tau) - \boldsymbol{R}(t) \rangle |_{t,N} \tag{4}$$

In (4), $\tau$ is the lag time. Next, we extracted the diffusion coefficient of the beads by fitting the MSD to the following equation:

$$MSD(\tau) = 4D\tau^{\alpha} + N \tag{5}$$

Here, $D$ is the diffusion coefficient, $\alpha$ is the diffusivity exponent, and $N$ is a term to account for the tracking noise. For fluids with terminally viscous behavior, $\alpha$ approaches 1 at long lag times which allows for the calculation of the terminal viscosity through the Stokes-Einstein equation:

$$\eta = \frac{k_B T}{6\pi D r} \tag{6}$$

Here, $r$ is the particle radius, $k_B$ is the Boltzmann coefficient, and $T$ is the temperature in Kelvin. The averaged value was reported for the viscosity from several condensates. The error was estimated as the standard deviation.

For the probability density analysis reported in **Fig. 3** of the main text, we computed the time averaged MSD of individual particles at a lag time of 10 seconds.

$$MSD_i(10) = \langle \boldsymbol{R_i}(t+10) - \boldsymbol{R_i}(t) \rangle |_t \tag{7}$$

The values for the root-mean square displacement (RMSD = $\sqrt{MSD}$) were then plotted as normalized histograms using the matplotlib python library.

For the calculation of the average diffusion coefficient, we split the long movie that we collected into shorter movies consisting of 200 frames (20 seconds) starting at different time points from the start of the experiment. For each movie, the apparent instantaneous diffusion coefficient was calculated using:

$$D = \left.\frac{MSD(\tau)}{4\tau}\right|_{\tau=10} \quad (8)$$

We used this method of estimating $D$ since fitting the time-dependent MSD with equation (5) yielded diffusivity exponent values $\alpha < 1$, which results in the parameter $D$ having units that are different from a canonical diffusion coefficient. Finally, we plotted the apparent average diffusion coefficients as a function of time.

**Flow activation energy analysis [11]**

Viscosities of A1-LCD condensates were calculated at different temperatures as per the description of the temperature-controlled VPT section. The viscosities are expected to decay exponentially with increasing temperature according to the Arrhenius law of viscosity [13]:

$$\eta = \eta_0 \exp\left(\frac{E_A}{RT}\right) \quad (9)$$

In (9), $\eta_0$ is the pre-exponential entropic factor, $E_A$ is the flow activation energy, $R$ is the universal gas constant, and $T$ is the temperature. Taking a natural logarithm of both sides of this equation yields

$$\ln \eta = \ln \eta_0 + \frac{E_A}{R}\left(\frac{1}{T}\right) \quad (10)$$

We plotted the natural log of viscosity against $1/T$ and fitted the data with a linear equation $y = mx + b$. The slope of the straight line is $m = \frac{E_A}{R}$, from which we calculated the flow activation energy [11]. These experiments were repeated three times, and the average value of the activation energy was reported. The error was taken as the standard deviation.

**Calculation of the complex shear modulus from video particle tracking microrheology of allY$^{30GtoS}$ condensates**

Since the motion of embedded particles within a fluid is driven by the thermal fluctuations of the fluid itself, the viscoelastic properties of the fluid can be estimated by analyzing the mean squared displacement (MSD) of the embedded probe particles. We do this by first calculating the fluid compliance from the MSD using [14]:

$$J(t) = \frac{3\pi a}{N_d k_B T} \langle \Delta r^2(\tau) \rangle \tag{11}$$

Here, $a$ is the particle radius, $N_d$ is the dimensionality of the probe particle motion ($N_d$=2 in our case), $k_B$ is the Boltzmann constant, and $T$ is the absolute temperature. Since the relaxation modulus $G(t)$ can be related to the compliance $J(t)$ by a convolution integral

$$\int_0^\tau G(t)J(\tau - t)dt = \tau, \tag{12}$$

the complex shear modulus can then be calculated using a Fourier transform as follows:

$$G^*(\omega) = \frac{1}{i\omega \hat{J}(\omega)} \tag{13}$$

where $\hat{J}(\omega)$ is the compliance in the Fourier space. We used two methods, one proposed by Evans et al., [15] and the other by Mason et al., [16] to convert the time-space compliance to frequency-space shear modulus $G^*(\omega)$.

In the method of Evans et al., the complex shear modulus $G^*(\omega)$ can be computed using:

$$\frac{i\omega}{G^*(\omega)} = i\omega J(0) + \frac{\left(1 - e^{-i\omega t_1}\right)\left(J_1 - J(0)\right)}{t_1} + \frac{e^{-i\omega t_N}}{\eta} + \sum_{k=2}^{N} \left(\frac{J_k - J_{k-1}}{t_k - t_{k-1}}\right)\left(e^{-i\omega t_{k-1}} - e^{-i\omega t_k}\right) \tag{14}$$

where $(t_i, J_i)$ are the discrete experimental data points of the calculated compliance. The advantage of using the method of Evans et al.,[15] is that there is no fitting required of the data. This distinguishes it from the method of Mason et al., [16] where power law fitting is needed to convert the MSD to the rheological moduli. To improve the calculation, we oversampled the MSD data from our VPT measurements on AllY$^{30GtoS}$ at different observation times (Main text **Fig. 3f**) using a cubic spline (**Supplementary Fig. S2**). The value of $J(0)$ was estimated by extrapolating the experimental data to $t = 0$ by a linear fit of the first four data points. Further, the value of $\eta$ was estimated by a linear fit of the last ten points (using $\eta = 1/\dot{J}(t)$).

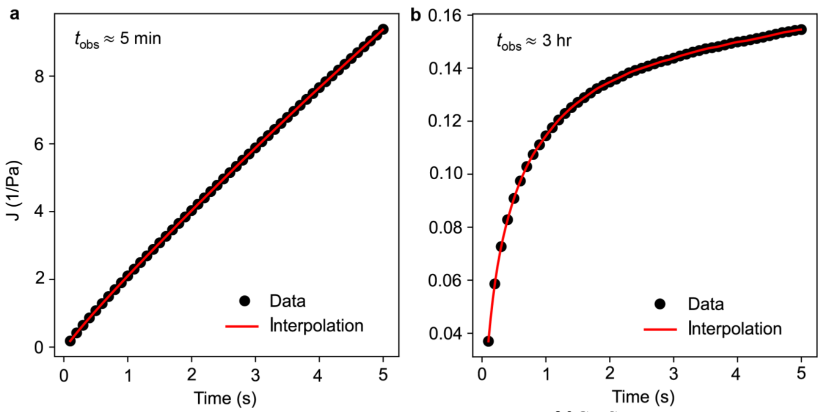


**Supplementary Figure S2.** Calculated compliance of allY$^{30GtoS}$ condensates at (**a**) $t_{obs} \approx 5$ min and (**b**) $t_{obs} \approx 3$ hrs (right). Also plotted are the oversampled compliance functions (red) using a cubic spline. The compliance is calculated directly from the MSD data shown in **Fig. 3f** in the main text using equation 11.

The computation of the complex shear modulus can also be independently performed using the method of Mason et al.[16], which involves approximating the creep compliance as a power function.

$$J(t) = J_0 t^{\alpha} \tag{15}$$

where $\alpha$ is the diffusivity parameter which provides information on the nature of the diffusion process with $\alpha = 1$ representing normal Brownian diffusion and $\alpha < 1$ signifying a sub-diffusive process.

To estimate the complex shear modulus $G^*(\omega)$, the interpolated creep compliance (**Supplementary Fig. S2**) above was fitted locally to a power function. The neighbors in the local fit were weighted using a Gaussian function as described previously [16]. The fitting provides $\alpha$ at $\omega = 1/t$ that can be used to compute the complex shear modulus at frequencies $\omega = 1/t$ using [17]

$$G^*(\omega) = \frac{e^{i\pi\alpha/2}}{J\left(t = \frac{1}{\omega}\right)\Gamma(1+\alpha)} \tag{16}$$

where Γ is the gamma function.

Upon converting compliance data to complex moduli, using the methods of Evans et al., [15] as well as Mason et al., [16] we observe that allY$^{30GtoS}$ condensates are dominantly viscous at $t_{obs} \approx$ 5 min (**Extended Data Fig. 5a-5c**). However, the elastic modulus of these condensates increases dramatically upon physical aging ($t_{obs} \approx 3$ hrs). The overall shapes of the frequency-dependent viscoelastic moduli of aged allY$^{30GtoS}$ condensates resemble that of Kelvin-Voigt solids (**Extended Data Figs. 5d, 5e**). Both methods yield consistent results (**Extended Data Fig. 5c, 5f**), confirming that allY$^{30GtoS}$ condensates are dominantly elastic at long observation times ($t_{obs} \geq 3$ hrs).

### Graph-theoretic implementation of Rouse-Zimm theory

In both the single-chain and collective models, all rheological properties were calculated from the eigenvalues $\lambda_p$ of the Zimm matrix and averaged over the ensemble. The first eigenvalue is zero [18]. For the $p^{\text{th}}$ mode, each non-zero eigenvalue is inversely proportional to the relaxation time $\tau_p$, given by [19]:

$$\tau_p = \frac{\zeta b^2}{6 k_B T \lambda_p} \tag{17}$$

In (17), $k_B$ is the Boltzmann constant and $T$ is the temperature. The Kuhn length $b$ and friction coefficient $\zeta$ of the medium are set to unity. To calculate the dynamical moduli, we first calculated the relaxation modulus $G(t)$ from a linear superposition of the eigenmodes [20] as a function of time $t$:

$$G(t) = \frac{\phi k_B T}{N b^3} \sum_{p=1}^{N-1} g_p e^{-t/\tau_p} \tag{18}$$

In (18), $N$ is the number of nodes (beads in the Rouse-Zimm theory). We set the weights $g_p$ and volume fraction $\phi$ to be unity. The latter choice reflects the fact the Rouse-Zimm theory was designed for polymer melts. This choice was also made to enable unbiased rescaling to put the computed moduli on the same footing as the experimental data.

The frequency-dependent complex shear modulus $G^*(\omega)$ was calculated from the Fourier transform of equation 18 [20]. The computed frequency range is greater than the shortest and longest relaxation times to ensure that we capture all the relevant dynamics. The values of $G^*(\omega)$ and $\omega$ at the single point corresponding to the crossover of the storage and loss moduli were rescaled by a multiplicative factor so the computed and measured crossover frequencies match one another. This represents a single-parameter-based rescaling that puts the computed and measured moduli on the same scale, thereby allowing us to quantitatively assess if the computed moduli match the measured moduli. Note that the computations only use information about the ensemble-averaged network structure within condensates. If information that is missing from the simulations ends up contributing to most of the measured moduli, then rescaling to match the crossover frequency, which uniformly shifts the computed frequency spectrum, will not lead to a matching of the computed and measured moduli. However, as shown in the main text, the simple rescaling operation is sufficient not only to bring the measured and computed moduli onto the same scale, but we also observe close correspondence across several orders of magnitudes of frequencies. We find that the agreement between computed and measured moduli is superior for the collective model, especially for computed storage moduli. This highlights the fact that storage moduli depend on the collective behavior of the condensate-spanning network. The use of the single, sequence-specific rescaling parameter is justified on the grounds that the weights $g_p$ in equation 18 are not known *a priori*. However, what we show is that we do not need to compute each of the weights separately. Instead, a single, uniform rescaling is sufficient for facilitating comparisons between experiments and computations.

Computations were carried out for all variants of A1-LCD reported in Farag et al [21], for FUS-LCD and for a homopolymer that was tuned to give rise to phase behavior closely matching that of A1-LCD WT$^{\text{-NLS}}$ [21] (**Table S2**).

Two other quantities, namely the zero-shear viscosity and the steady-state compliance, can be derived from the moments of the gyration tensor [22]. Using the spectrum of relaxation times given by equation 17, we can express the zero-shear viscosity as:

$$\eta_0 = \frac{\phi kT}{Nb^3} \sum_{p=1}^{N-1} \tau_p \tag{19}$$

The steady-state compliance is computed using:

$$J_0 = \frac{Nb^3}{\phi kT} \frac{\sum_{p=1}^{N-1} \tau_p^2}{\left(\sum_{p=1}^{N-1} \tau_p\right)^2} \tag{20}$$

where the volume fraction at a given temperature is computed using the volume fractions of the dense and dilute phases using: $\phi = \phi_{\text{dense}}/\phi_{\text{sat}}$.

**DIC Microscopy and ThT assay**

Differential interference contrast microscopy (DIC) and ThT fluorescent images were obtained at room temperature using a Zeiss 780 LSM confocal microscope with a 20× objective. Samples were prepared as described above. Protein concentrations were selected to ensure they were above the $c_{\text{sat}}$ of the corresponding variant at room temperature. 4 μL of the protein solution with 20 μM of ThT was applied to the slide and imaged at the indicated times. The ThT was excited with a 480 nm laser.

**Fourier Transform Infrared (FTIR) Spectroscopy**

The FTIR data were collected using a Nicolet iS20 FTIR instrument (ThermoFisher Scientific) with a Smart iTX Diamond accessory. For dense phase measurements phase separation was induced in 200 to 500 μL of 1 to 2 mM stock solutions of protein in 20 mM HEPES pH 7 by adding NaCl to a final concentration of 150 mM. The samples were then centrifuged at 5000 RCF for 5 min. A 3 μL sample of dense phase was transferred to the stage using a positive displacement pipette. The dilute phase was then pipetted around the dense phase and the sample was sealed with the liquid chamber accessory to prevent desiccation. The insulin fibrils were centrifuged and washed with deionized water 5 times, and then transferred to the stage for analysis. Each FTIR spectrum represents the average of 64 consecutive scans, with a resolution of 0.2411 $cm^{-1}$. A blank of 20 mM HEPES pH 7 with 150 mM NaCl using the same scan and resolution settings was subtracted from each spectrum. The spectra were checked for an appropriate ratio of amide I and II band intensity, low noise, and the absence of other artifacts to confirm proper data acquisition and background subtraction. Data were analyzed using OriginPro 2023 (OriginLab). The inverse second-derivative of the absorbance spectra in the amide I region was calculated using a third order Savitzky–Golay filter [23] with a 100–120-point window, depending on the data set. The inverse second-derivative was baseline-corrected assuming flat baseline segments connecting minima between 1710 and 1595 $cm^{-1}$. The curves for each sample were then fit using the same starting parameters of 7 Gaussians centered at 1614, 1632, 1643, 1653, 1667, 1680, and 1692 $cm^{-1}$, corresponding to aggregated β-strands, β-sheets, random coil, α-helix, $3_{10}$ helix, β-turn, and parallel β-sheet structures, respectively. The insulin fibrils were fit with peaks starting at 1625, 1667, and 1680 $cm^{-1}$, corresponding to aggregated β-strands, $3_{10}$ helix, β-turn, respectively.

**Insulin fibril preparation**

To generate the insulin fibril samples, lyophilized bovine insulin (Cell Applications 128100) was dissolved in 20% acetic acid and the concentration was adjusted to 2 mM. The samples were then incubated at 68°C overnight as previously described [24, 25]. For microscopy, the slide was assembled as described above, using double-sided tape, and sealing of the sample with mineral oil. The slide was then incubated on top of a heat block set to 75°C using Eppendorf lids as offsets to prevent direct contact with the block. The slide temperature was measured at 70°C using an infrared thermometer. The slide was incubated in the dark overnight.

**Deep Etch Electron Microscopy**

Samples for freeze fracture were prepared by exchanging into 20 mM HEPES pH 7 as described above. The concentration of allY$^{30GtoS}$ was adjusted for the final concentration to be 400-600 µM. NaCl was added to a final concentration of 150 mM to induce phase separation. The sample was then applied to a mica disk and either flash frozen immediately in liquid nitrogen or incubated overnight in a vapor chamber to allow the sample to age before flash freezing in nitrogen.

Replicas of deep etched samples for TEM were produced using a Leica EM ACE900 Freeze Fracture System. In brief, aged mica sandwiches were frozen by plunging into liquid nitrogen, separated beneath the surface of liquid nitrogen, and transferred to the ACE900 Freeze Fracture System. Then, water was sublimated at -80 °C for 1 hr, prior to shadowing with 7 nm Pt/C at an angle of 24°. The replica was supported by 8 nm carbon deposition at 85° before being washed through several DI water rinses and transferred to a copper TEM grid for TEM imaging (JEOL JEM-1400 Plus) at 120 kV.

**Software**

pMOT and VPT analysis were done using custom-made python scripts that are available on GitHub (see https://github.com/BanerjeeLab-repertoire/Material-properties). Fiji-ImageJ (version 1.53c) was used for image processing. Origin (2023) was used for Graphing. Adobe Illustrator CC (2019, v23.0) was used for the figure assembly and production. ZEN (blue, v2.3) and micromanager v2.0 were used for image collection on Zeiss Primovert microscopes. Bluelake (v1.6.11) was used for image recording and particle tracking using a Lumicks C-Trap microscope. FTIR spectra were processed and analyzed using OriginPro 2023. All details for reproducing the graph-theoretic analysis and custom-made scripts are available on GitHub (see: https://github.com/Pappulab/Material-properties).

### Supplementary Tables

**Table S1: Amino acid sequences of the variants used in pMOT and VPT measurements.**

| [a] | **Protein sequence[b]** | [c] |
|---|---|---|
| allF | GSMASASSSQRGRSGSGNFGGGRGGGFGGNDNFGRGGNFSGRGGFGGSRGGGGFGGS<br>GDGFNGFGNDGSNFGGGGSFNDFGNFNNQSSNFGPMKGGNFGGRSSGGSGGGGQFFA<br>KPRNQGGFGGSSSSSSFGSGRRF | 19F |
| WT | GSMASASSSQRGRSGSGNFGGGRGGGFGGNDNFGRGGNFSGRGGFGGSRGGGGYGGS<br>GDGYNGFGNDGSNFGGGGSYNDFGNYNNQSSNFGPMKGGNFGGRSSGPYGGGGQYFA<br>KPRNQGGYGGSSSSSSYGSGRRF | 8Y 12F |
| allY | GSMASASSSQRGRSGSGNYGGGRGGGYGGNDNYGRGGNYSGRGGYGGSRGGGGYGGS<br>GDGYNGYGNDGSNYGGGGSYNDYGNYNNQSSNYGPMKGGNYGGRSSGGSGGGGQYYA<br>KPRNQGGYGGSSSSSSYGSGRRY | 19Y |
| W- | GSMASASSSQRGRSGSGNSGGGRGGGWGGNDNWGRGGNSSGRGGWGGSRGGGGWGGS<br>GDGWNGWGNDGSNSGGGGSSNDWGNWNNQSSNWGPMKGGNWGGRSSGGSGGGGQWSA<br>KPRNQGGWGGSSSSSSSGSGRRW | 13W |
| YtoW | GSMASASSSQRGRSGSGNFGGGRGGGFGGNDNFGRGGNFSGRGGFGGSRGGGGWGGS<br>GDGWNGFGNDGSNFGGGGSWNDFGNWNNQSSNFGPMKGGNFGGRSSGSGGGGGQWFA<br>KPRNQGGWGGSSSSSSWGSGRRF | 7W 12F |
| FtoW | GSMASASSSQRGRSGSGNWGGGRGGGWGGNDNWGRGGNWSGRGGWGGSRGGGGYGGS<br>GDGYNGWGNDGSNWGGGGSYNDWGNYNNQSSNWGPMKGGNWGGRSSGSGGGGGQYWA<br>KPRNQGGYGGSSSSSSYGSGRRW | 12W 7Y |
| allW | GSMASASSSQRGRSGSGNWGGGRGGGWGGNDNWGRGGNWSGRGGWGGSRGGGGWGGS<br>GDGWNGWGNDGSNWGGGGSWNDWGNWNNQSSNWGPMKGGNWGGRSSGSGGGGGQWWA<br>KPRNQGGWGGSSSSSSWGSGRRW | 19W |
| WT$^{20GtoS}$ | GSMASASSSQRSRSGSGNFSGSRSGSFSGNDNFGRSGNFSGRSGFGGSRSGGGYSGS<br>GDSYNSFGNDGSNFSGSGSYNDFGNYNNQSSNFGPMKSGNFGGRSSGSSGGSGQYFA<br>KPRNQGSYSGSSSSSSYGSSRRF | 7Y 12F |
| WT$^{30GtoS}$ | GSMASASSSQRSRSSSGNFSGSRSGSFSGNDNFGRSGNFSGRSGFSGSRSGSGYSGS<br>SDSYNSFGNDSSNFSGSSSYNDFGNYNNQSSNFGPMKSGNFSGRSSSSSGSSGQYFA<br>KPRNQGSYSGSSSSSSYSSSRRF | 7Y 12F |
| allY$^{20GtoS}$ | GSMASASSSQRSRSGSGNYSGSRSGSYSGNDNYGRSGNYSGRSGYGGSRSGGGYSGS<br>GDSYNSYGNDGSNYSGSGSYNDYGNYNNQSSNYGPMKSGNYGGRSSGSSGGSGQYYA<br>KPRNQGSYSGSSSSSSYGSSRRY | 19Y |
| allY$^{30GtoS}$ | GSMASASSSQRSRSSSGNYSGSRSGSYSGNDNYGRSGNYSGRSGYSGSRSGSGYSGS<br>SDSYNSYGNDSSNYSGSSSYNDYGNYNNQSSNYGPMKSGNYSGRSSSSSGSSGQYYA<br>KPRNQGSYSGSSSSSSYSSSRRY | 19Y |

[a] variant name; [b] the N-terminal GS-extensions are leftovers of TEV cleavage sites; [c] number and type of aromatic residues in the sequence

**Table S2: Amino acid sequences of the variants for which computations were performed and reported in Fig. 2a**. The simulation results for these variants were taken from the work of Farag et al [21]. Following this work, simulations of FUS-LCD and of a homopolymer whose properties were tuned to give rise to phase behavior closely matching that of A1-LCD WT$^{-NLS}$ [21] were also analyzed.

| Construct | Amino acid sequence |
|---|---|
| WT$^{-NLS}$ | GSMASASSSQRGRSGSGNFGGGRGGGFGGNDNFGRGGNFSGRGGFGGSRGGGGYGGSGDGY<br>NGFGNDGSNFGGGGSYNDFGNYNNQSSNFGPMKGGNFGGRSSGGSGGGGQYFAKPRNQGGY<br>GGSSSSSSYGSGRRF |
| WT$^{+NLS}$ | GSMASASSSQRGRSGSGNFGGGRGGGFGGNDNFGRGGNFSGRGGFGGSRGGGGYGGSGDGY<br>NGFGNDGSNFGGGGSYNDFGNYNNQSSNFGPMKGGNFGGRSSGPYGGGGQYFAKPRNQGGY<br>GGSSSSSSYGSGRRF |
| -12F+12Y (allY) | GSMASASSSQRGRSGSGNYGGGRGGGYGGNDNYGRGGNYSGRGGYGGSRGGGGYGGSGDGY<br>NGYGNDGSNYGGGGSYNDYGNYNNQSSNYGPMKGGNYGGRSSGGSGGGGQYYAKPRNQGGY<br>GGSSSSSSYGSGRRY |
| +7F-7Y (allF) | GSMASASSSQRGRSGSGNFGGGRGGGFGGNDNFGRGGNFSGRGGFGGSRGGGGFGGSGDGF<br>NGFGNDGSNFGGGGSFNDFGNFNNQSSNFGPMKGGNFGGRSSGGSGGGGQFFAKPRNQGGF<br>GGSSSSSSFGSGRRF |
| -4F-2Y | GSMASASSSQRGRSGSGNSGGGRGGGFGGNDNFGRGGNSSGRGGFGGSRGGGGYGGSGDGY<br>NGFGNDGSNSGGGGSSNDFGNYNNQSSNFGPMKGGNFGGRSSGGSGGGGQYSAKPRNQGGY<br>GGSSSSSSSGSGRRF |
| -9F+6Y | GSMASASSSQRGRSGSGNFGGGRGGGYGGNDNYGRGGNYSGRGGFGGSRGGGGYGGSGDGY<br>NGGGNDGSNYGGGGSYNDSGNYNNQSSNFGPMKGGNYGGRSSGGSGGGGQYGAKPRNQGGY<br>GGSSSSSSYGSGRRY |
| -8F+4Y | GSMASASSSQRGRSGSGNFGGGRGGGYGGNDNGGRGGNYSGRGGFGGSRGGGGYGGSGDGY<br>NGGGNDGSNYGGGGSYNDSGNYNNQSSNFGPMKGGNYGGRSSGGSGGGGQYGAKPRNQGGY<br>GGSSSSSSYGSGRRF |
| -9F+3Y | GSMASASSSQRGRSGSGNFGGGRGGGYGGNDNGGRGGNYSGRGGFGGSRGGGGYGGSGDGY<br>NGGGNDGSNYGGGGSYNDSGNGNNQSSNFGPMKGGNYGGRSSGGSGGGGQYGAKPRNQGGY<br>GGSSSSSSYGSGRRS |
| +12D | GSMASADSSQRDRDDSGNFGDGRGGGFGGNDNFGRGGNFSDRGGFGGSRGDGGYGGDGDGY<br>NGFGNDGSNFGGGGSYNDFGNYNNQSSNFDPMKGGNFGDRSSGPYDGGGQYFAKPRNQGGY<br>GGSSSSSSYGSDRRF |
| +8D | GSMASASSSQRDRSGSGNFGGGRDGGFGGNDNFGRGDNFSGRGDFGGSRDGGGYGGSGDGY<br>NGFGNDGSNFGGGGSYNDFGNYNNQSSNFGPMKGGNFGGRSSDPYGGGGQYFAKPRNQDGY<br>GGSSSSSSYDSGRRF |

| | |
|---|---|
| +4D | GSMASASSSQRDRSGSGNFGGGRGGGFGGNDNFGRGGNFSGRGDFGGSRGGGGYGGSGDGY<br>NGFGNDGSNFGGGGSYNDFGNYNNQSSNFGPMKGGNFGGRSSDPYGGGGQYFAKPRNQGGY<br>GGSSSSSSYDSGRRF |
| -4D | GSMASASSSQRGRSGSGNFGGGRGGGFGGNGNFGRGGNFSGRGGFGGSRGGGGYGGSGGGY<br>NGFGNSGSNFGGGGSYNGFGNYNNQSSNFGPMKGGNFGGRSSGPYGGGGQYFAKPRNQGGY<br>GGSSSSSSYGSGRRF |
| +2R | GSMASASSSQRGRSGSGNFGGGRGGGFGGNDNFGRGGNFSGRGGFGGSRGGGGYGGSGDGY<br>NGFRNDGSNFGGGGRYNDFGNYNNQSSNFGPMKGGNFGGRSSGPYGGGGQYFAKPRNQGGY<br>GGSSSSSSYGSGRRF |
| -6R | GSMASASSSQGGRSGSGNFGGGRGGGFGGNDNFGGGGNFSGSGGFGGSRGGGGYGGSGDGY<br>NGFGNDGSNFGGGGSYNDFGNYNNQSSNFGPMKGGNFGGSSSGPYGGGGQYFAKPGNQGGY<br>GGSSSSSSYGSGGRF |
| -10R | GSMASASSSQGGSSGSGNFGGGGGGGFGGNDNFGGGGNFSGSGGFGGSGGGGGYGGSGDGY<br>NGFGNDGSNFGGGGSYNDFGNYNNQSSNFGPMKGGNFGGSSSGPYGGGGQYFAKPGNQGGY<br>GGSSSSSSYGSGGGF |
| -3R+3K | GSMASASSSQRGKSGSGNFGGGRGGGFGGNDNFGRGGNFSGRGGFGGSKGGGGYGGSGDGY<br>NGFGNDGSNFGGGGSYNDFGNYNNQSSNFGPMKGGNFGGRSSGGSGGGGQYFAKPRNQGGY<br>GGSSSSSSYGSGRKF |
| -6R+6K | GSMASASSSQKGKSGSGNFGGGRGGGFGGNDNFGKGGNFSGRGGFGGSKGGGGYGGSGDGY<br>NGFGNDGSNFGGGGSYNDFGNYNNQSSNFGPMKGGNFGGKSSGGSGGGGQYFAKPRNQGGY<br>GGSSSSSSYGSGRKF |
| +7K+12D | GSMASADSSQRDRDDKGNFGDGRGGGFGGNDNFGRGGNFSDRGGFGGSRGDGKYGGDGDKY<br>NGFGNDGKNFGGGGSYNDFGNYNNQSSNFDPMKGGNFKDRSSGPYDKGGQYFAKPRNQGGY<br>GGSSSSKSYGSDRRF |
| -20G+20S<br>(WT$^{20GtoS}$) | GSMASASSSQRSRSGSGNFSGSRSGSFSGNDNFGRSGNFSGRSGFGGSRSGGGYSGSGDSY<br>NSFGNDGSNFSGSGSYNDFGNYNNQSSNFGPMKSGNFGGRSSGSSGGSGQYFAKPRNQGSY<br>SGSSSSSSYGSSRRF |
| -30G+30S<br>(WT$^{30GtoS}$) | GSMASASSSQRSRSSSGNFSGSRSGSFSGNDNFGRSGNFSGRSGFSGSRSGSGYSGSSDSY<br>NSFGNDSSNFSGSSSYNDFGNYNNQSSNFGPMKSGNFSGRSSSSSGSSGQYFAKPRNQGSY<br>SGSSSSSSYSSSRRF |
| -12F+12Y-20G+20S<br>(allY$^{20GtoS}$) | GSMASASSSQRSRSGSGNYSGSRSGSYSGNDNYGRSGNYSGRSGYGGSRSGGGYSGSGDSY<br>NSYGNDGSNYSGSGSYNDYGNYNNQSSNYGPMKSGNYGGRSSGSSGGSGQYYAKPRNQGSY<br>SGSSSSSSYGSSRRY |
| -12F+12Y-30G+30S<br>(allY$^{30GtoS}$) | GSMASASSSQRSRSSSGNYSGSRSGSYSGNDNYGRSGNYSGRSGYSGSRSGSGYSGSSDSY<br>NSYGNDSSNYSGSSSYNDYGNYNNQSSNYGPMKSGNYSGRSSSSSGSSGQYYAKPRNQGSY<br>SGSSSSSSYSSSRRY |

| | |
|---|---|
| +7F-7Y-20G+20S (allF$^{20GtoS}$) | GSMASASSSQRSRSGSGNFSGSRSGSFSGNDNFGRSGNFSGRSGFGGSRSGGGFSGSGDSF<br>NSFGNDGSNFSGSGSFNDFGNFNNQSSNFGPMKSGNFGGRSSGSSGGSGQFFAKPRNQGSF<br>SGSSSSSSFGSSRRF |
| +7F-7Y -30G+30S (allF$^{30GtoS}$) | GSMASASSSQRSRSSSGNFSGSRSGSFSGNDNFGRSGNFSGRSGFSGSRSGSGFSGSSDSF<br>NSFGNDSSNFSGSSSFNDFGNFNNQSSNFGPMKSGNFSGRSSSSSGSSGQFFAKPRNQGSF<br>SGSSSSSSFSSSRRF |
| -23S+23T | GSMATATTTQRGRTGTGNFGGGRGGGFGGNDNFGRGGNFTGRGGFGGTRGGGGYGGTGDGY<br>NGFGNDGTNFGGGGTYNDFGNYNNQTTNFGPMKGGNFGGRTTGGTGGGGQYFAKPRNQGGY<br>GGTTTTTTYGTGRRF |
| -14N+14Q | GSMASASSSQRGRSGSGQFGGGRGGGFGGQDQFGRGGQFSGRGGFGGSRGGGGYGGSGDGY<br>QGFGQDGSQFGGGGSYQDFGQYQQQSSQFGPMKGGQFGGRSSGGSGGGGQYFAKPRQQGGY<br>GGSSSSSSYGSGRRF |
| -14N-4Q+18G | GSMASASSSGRGRSGSGGFGGGRGGGFGGGDGFGRGGGFSGRGGFGGSRGGGGYGGSGDGY<br>GGFGGDGSGFGGGGSYGDFGGYGGGSSGFGPMKGGGFGGRSSGGSGGGGGYFAKPRGGGGY<br>GGSSSSSSYGSGRRF |
| +7R+10D | GSMASADSSQRDRDGRGNFGDGRGGGFGGNDNFGRGGNFSDRGGFGGSRGGGRYGGDGDRY<br>NGFGNDGRNFGGGGSYNDFGNYNNQSSNFDPMKGGNFRDRSSGPYDRGGQYFAKPRNQGGY<br>GGSSSSRSYGSDRRF |
| +7R+12D | GSMASADSSQRDRDDRGNFGDGRGGGFGGNDNFGRGGNFSDRGGFGGSRGDGRYGGDGDRY<br>NGFGNDGRNFGGGGSYNDFGNYNNQSSNFDPMKGGNFRDRSSGPYDRGGQYFAKPRNQGGY<br>GGSSSSRSYGSDRRF |
| -10F +7R+12D | GSMASADSSQRDRDDRGNGGDGRGGGGGGNDNGGRGGNGSDRGGGGGSRGDGRYGGDGDRY<br>NGGGNDGRNGGGGGSYNDGGNYNNQSSNGDPMKGGNGRDRSSGPYDRGGQYGAKPRNQGGY<br>GGSSSSRSYGSDRRG |
| -12F+12Y-10R | GSMASASSSQGGSSGSGNYGGGGGGGYGGNDNYGGGGNYSGSGGYGGSGGGGGYGGSGDGY<br>NGYGNDGSNYGGGGSYNDYGNYNNQSSNYGPMKGGNYGGSSSGPYGGGGQYYAKPGNQGGY<br>GGSSSSSSYGSGGGY |
| -SYNDFG | GSMASASSSQRGRSGSGNFGGGRGGGFGGNDNFGRGGNFSGRGGFGGSRGGGGYGGSGDGY<br>NGFGNDGSNFGGGGNYNNQSSNFGPMKGGNFGGRSSGGSGGGGQYFAKPRNQGGYGGSSSS<br>SSYGSGRRF |
| FUS-LCD | GSMASNDYTQQATQSYGAYPTQPGQGYSQQSSQPYGQQSYSGYSQSTDTSGYGQSSYSSYG<br>QSQNTGYGTQSTPQGYGSTGGYGSSQSSQSSYGQQSSYPGYGQQPAPSSTSGSYGSSSQSS<br>SYGQPQSGSYSQQPSYGGQQQSYGQQQSYNPPQGYGQQNQYNSSSGGGGGGGGGGNYGQDQ<br>SSMSSGGGSGGGYGNQDQSGGGGSGGYGQQDRG |

**Supplementary Movie legends**

**Supplemental video 1.** Representative video of a WT$^{30GtoS}$ condensate undergoing physical aging as seen by the arrest of 200 nm probe particle dynamics within the condensate.

**Supplemental video 2.** Representative video of a creep test for an allY condensate at 24 hours after sample preparation. The bead diameter is 2 μm.

**Supplemental video 3.** Representative video of a creep test for an allY$^{20GtoS}$ condensate at 24 hours after sample preparation. The bead diameter is 2 μm.
